# Single-cell Tree-based Model for Genomic-Disease Association

**DOI:** 10.64898/2025.12.31.697253

**Authors:** Zhikang Liu, Yiyang Niu, Tian Le, Daniel G Chen, Yapeng Su, Ye Zheng

**Affiliations:** Department of Bioinformatics and Computational Biology, The University of Texas MD Anderson Cancer Center, Houston, TX, USA; Program in Immunology, Clincal Research Division and Translational Science and Therapeutics Division, Fred Hutchinson Cancer Center, Seattle, WA, USA; Department of Systems Biology, The University of Texas MD Anderson Cancer Center, Houston, TX, USA

## Abstract

The rapid maturation of single-cell multi-omics technologies has enabled unprecedented resolution for mapping disease states and identifying disease-associated biomarkers. In practice, biomarkers are often discovered through differential detection that treat genomic features as independent contributors to phenotypes, while the combinatorial interactions that drive clinical outcomes remain a practical challenge. We present scanCT (single-cell analysis of Clinical Tree), a tree-based framework that identifies groups of genomic features associated with distinct disease phenotypes in a highly interpretable manner. scanCT uses an unbiased, model-based variable-selection procedure for data-driven split selection, which is important for handling the diverse distributional properties of single-cell data across modalities. The tree architecture captures feature interaction effects, and the association modeling enables adjustment for confounding factors. We apply scanCT to longitudinal single-cell multi-omics COVID-19 datasets spanning diverse clinical outcomes and multiple time points per patient. scanCT identifies phenotype-specific gene and protein markers while accounting for age and sex, and it reveals interpretable synergistic marker combinations that help explain differences in patient clinical phenotypes.

## Introduction

Unraveling the molecular determinants of complex disease phenotypes is a central goal of precision medicine ^1^. The advance of single-cell multi-omics technologies enables joint profiling of gene expression, surface protein abundance ^2^, chromatin accessible ^3^, methylation and three-dimensional chromatin structure ^4^ at unprecedented resolution, providing us with high dimensional biomarkers to investigate the genomic-disease association ^5^.

A key analytical challenge in the association of genomic features with disease is to identify molecular features and reveal their joint effects contributing to clinical outcomes. Tree-based models are well-suited to this setting because they naturally capture non-linear effects and higher-order interactions without assuming independency across features or linear relationship with the phenotypes ^6^. In particular, tree-based models offer a distinct advantage in clinical applications in terms of high interpretability. They provide an intrinsically representation of the decision logic through a set of transparent hierarchical rules. The most widely adopted framework for this purpose is Classification and Regression Trees (CART)^6^, which employs a recursive partitioning strategy to segment patients or cells into risk groups. However, despite its interpretability, the standard greedy search algorithm used in CART introduces a critical limitation regarding how features are prioritized during model construction. Ensemble methods such as Random Forests and gradient-boosted trees (e.g., XGBoost) ^7,8^ are also widely used. Although these ensembles often achieve strong predictive performance, the feature interactions and interpretability are lost going from tree-based to a forest-based structure. Their variable-importance measures and selected split variables can be systematically distorted by variable selection bias. A well-documented source of bias arises from greedy split selection, a fundamental characteristic of standard recursive partitioning that is also inherited by many ensemble methods ^6,9^. These algorithms typically identify the optimal split by performing an exhaustive search over all candidate cut-points to maximize impurity reduction ^10^. As a consequence, predictors with many potential cut points, typically continuous gene-expression variables, are preferentially selected compared to variables with fewer unique values, such as binary or categorical protein markers. Even if protein markers carry strong and unique biological signals^9^, they tend not to be selected compared to gene features with many more unique values. Such a cardinality-driven biase can yield spurious associations, mask true drivers, and undermine the reliability and interpretability of feature-importance rankings ^11^. In multi-omics settings, where modalities differ sharply in measurement scale and discretization, these biases can be particularly problematic for biomarker prioritization.

To address the practical challenges, we developed scanCT (single-cell analysis of Clinical Trees, Fig. 1), an interpretable tree-based framework built on the GUIDE (Generalized, Unbiased, Interaction Detection and Estimation) family of algorithms^12,13^. GUIDE decouples split-variable selection from split-point optimization by using chi-squared hypothesis testing for variable screening, thereby mitigating selection bias toward high-cardinality predictors while retaining the ability to detect interactions. In addition, GUIDE logistic regression trees allow confounders to be incorporated as covariates within terminal-node models, enabling robust risk estimation and interpretable adjustment for demographic effects.

**Fig. 1.**
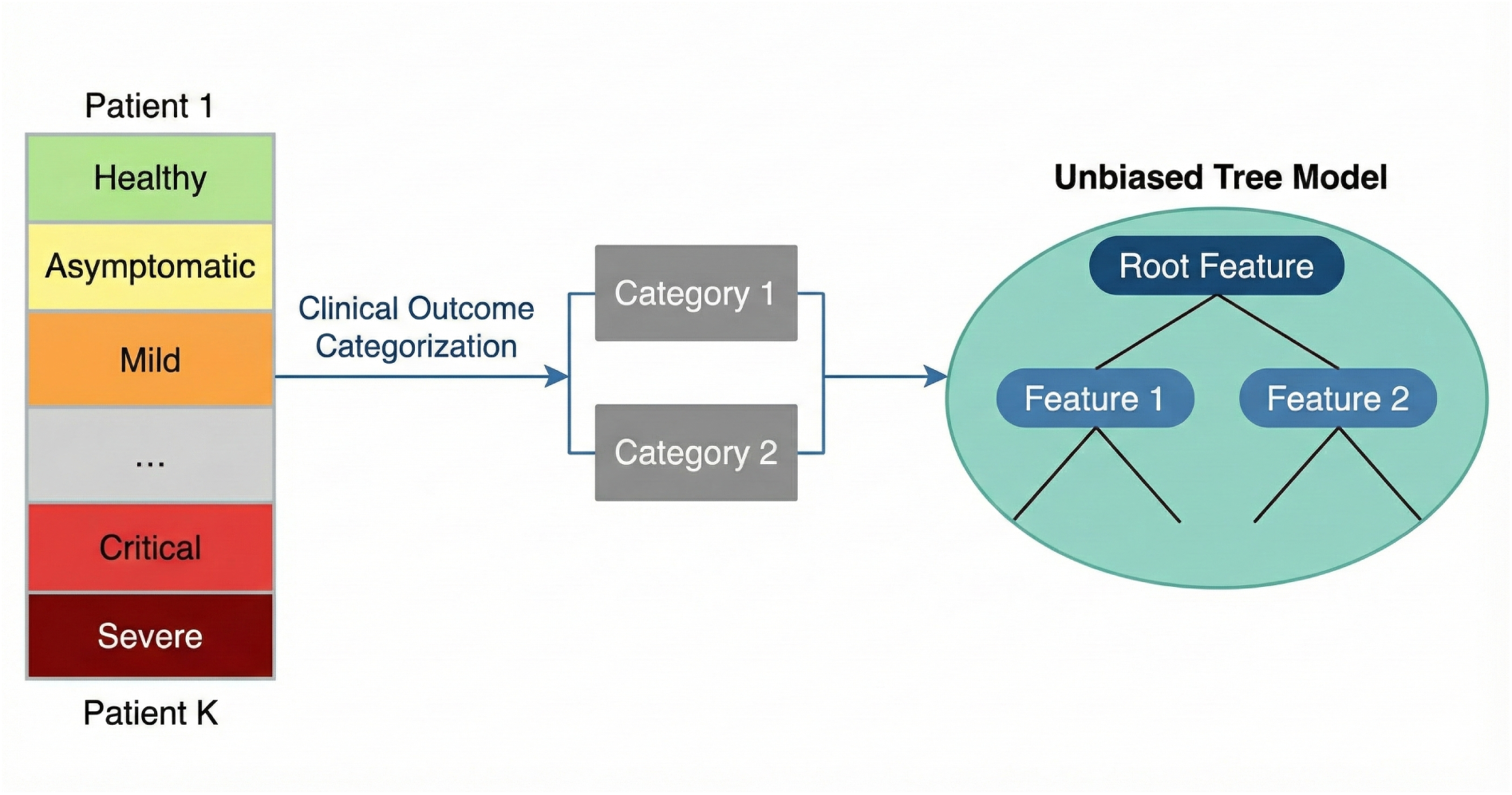
Overview of the scanCT analytical framework. Patient clinical statuses, ranging from healthy to severe, are consolidated into different categories through clinical outcome categorization. These defined endpoints serve as the response variable for the unbiased decision tree model, which is employed to identify robust genomic associations.

Illustrated through a series of COVID-19 studies ^14–16^, we characterize feature selection bias in conventional tree-based learners on single-cell multi-omics data. We demonstrate that scanCT yields importance scores and selected predictors that are less sensitive to predictor cardinality and correlation structure. Additionally, we benchmark scanCT baselines to quantify the performance and extend the framework to confounder-adjusted risk modeling using logistic regression trees. Together, these analyses provide a practical path toward bias-free, interpretable biomarker discovery and clinical risk stratification in single-cell multi-omics studies.

## Results

### Characterization of feature selection bias in single-cell multi-omics data

To characterize feature selection bias in tree-based learning on single-cell multi-omics data, we analyzed *N* = 110 patients with an average of 5,000 cells per patient and 2,587 features (2,500 highly variable genes and 87 protein markers with bimodal pattern) per cell. Predictors include continuous gene expression and binarized protein abundances measured by antibody-derived tags (ADT) from the CITE-seq data, ^17^ yielding a heterogeneous feature space with contrasting sparsity and cardinality and well-suited to probing selection bias. COVID-19 severity was encoded on an ordinal scale from 0 (healthy) to 5 (critical) (Methods). Based on this scale, we defined four comparison scenarios (G1–G4) to span broad case–control discrimination and fine-grained progression (Table 1). Cells were stratified into major immune lineages (Mononuclear phagocytes and HSPCs, B cells, CD4T cells, CD8T cells and T cells) based on the annotations provided from the original publication (Supplementary Table S1).^14^

**Table 1.** Summary of Data Grouping Based on Severity Levels.

| Group | Description |
| --- | --- |
| G1 | Healthy (0) vs Non-healthy or COVID (1) |
| G2 | Healthy (0) vs Asymptomatic + Mild (1) |
| G3 | Healthy (0) vs Critical + Severe (1) |
| G4 | Healthy (0) vs Asymptomatic + Mild (1) vs Critical + Severe (2) |

To assess whether scanCT selects splitting variables without favoring high-cardinality predictors, we conducted a permutation experiment on Mononuclear cells in Group 4 (Healthy vs Asymptomatic/Mild vs Severe/Critical). In each of 20 replicates, the response *Y* was randomly permuted to break any association with the predictors, creating a null setting in which all features are approximately independent of the outcome. Under this null, an unbiased method should show no systematic relationship between the number of unique values and the resulting importance scores.

We found the fitted regression lines show strong positive correlations between feature cardinality and importance in CART and RF with Pearson correlation coefficients of *r*_CART_ = 0.769 and *r*_RF_ = 0.859 (Fig. 2A, B).^7^ The top 20 ranked features (red points) are almost exclusively high-cardinality genes, such as S100A8 and CD74, each with more than 50,000 distinct values. These patterns indicate that, even under a null model with no signal, CART and RF systematically concentrate importance on high-cardinality predictors. By contrast, the scanCT shows a nearly horizontal regression line with negligible correlation between cardinality and importance with *r*_scanCT_ = 0.005 (Fig. 2C). The top-ranked features span a broad range of unique-value counts rather than clustering in the high-cardinality regime, and their error bars overlap extensively across the cardinality axis. Together, these observations demonstrate that scanCT maintains structural neutrality in variable selection under the null, assigning importance independently of feature cardinality and thereby avoiding the spurious preference for highly granular predictors observed in CART and RF.

**Fig. 2.**
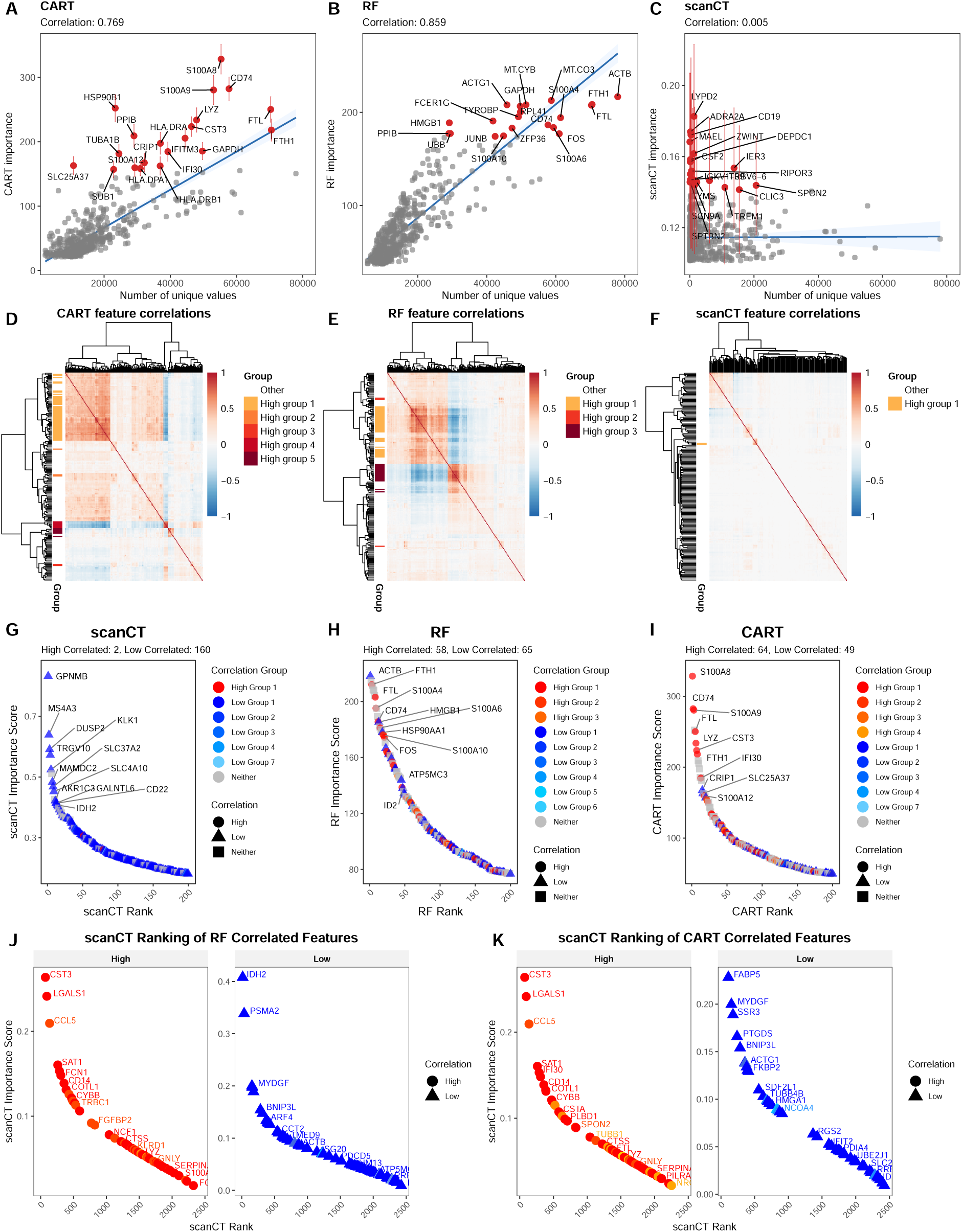
Characterization of feature selection bias in tree-based models. (A-C) Scatter plots of feature importance versus number of unique values for CART (A), Random Forest (RF; B), and scanCT (C) from a permutation experiment on Mononuclear Group 4 (20 null replicates with *Y* independent of *X*). Blue lines show fitted linear regressions with Pearson correlation coefficients; red points denote the top 20 ranked features with error bars indicating the standard deviation of importance across permutations. **(D–F)** Pairwise Pearson correlation heatmaps for the top 200 features selected by CART (D), RF (E), and scanCT (F) in Mononuclear Group 4, ordered by hierarchical clustering. CART and RF show dense, blockdiagonal clusters of strongly correlated predictors, whereas scanCT yields a more diffuse pattern with fewer large, highly correlated blocks. **(G–I)** Rank–abundance plots for the top 200 features selected by CART (G), RF (H), and scanCT (I), stratified by correlation status (highly correlated, low correlated, neither). Highly correlated predictors dominate the upper ranks for CART and RF, whereas low- and uncorrelated features primarily occupy scanCT’s top positions. **(J–K)** Cross-method comparison of scanCT rankings for features belonging to high-and low-correlation groups defined by CART (J) and RF (K). High-correlation groups from CART and RF are dispersed across the scanCT ranking rather than clustering at the top, indicating that scanCT does not inherit the collinearity bias of the comparison methods.

Additionally, in our analysis of the real clinical data, we examined the top 50 features identified by each method by ranking importance scores on CD8T G1. We observed that while standard ensemble methods (RF and XGBoost) ^7,8^ systematically demoted protein (ADT) features in favor of high-cardinality gene expression features, scanCT identified a distinct distribution of top-ranked features, capturing numerous surface protein markers that were effectively masked in the ensemble tree rankings (Supplementary Fig. S1). This suggests that scanCT’s unbiased selection strategy is particularly advantageous for detecting biologically relevant signals, such as surface proteins, that may otherwise be overshadowed by the high cardinality of transcriptomic data.

The scanCT algorithm is also designed to mitigate selection bias toward clusters of highly correlated features. In contrast, CART and RF tend to repeatedly select collinear predictors, thereby inflating their importance and reducing the effective dimensionality of the model ^18,19^.

To evaluate correlation bias, we computed the pairwise Pearson correlation matrix over the entire feature space. We defined highly correlated groups as clusters with *ρ >* 0.6 and lowly correlated groups as those with *ρ <* 0.1. For each method, we then extracted the top 200 features ranked by importance scores and examined whether correlated clusters disproportionately occupied the highest ranks.

We generated correlation heatmaps for the top 200 features on Mononuclear Group 4, ordered by hierarchical clustering. We noticed that CART and RF exhibited dense, block-diagonal structures of strong positive correlations (Fig. 2D-E), indicating that their top-ranked predictors are dominated by tightly collinear groups and high-cardinality, inter-correlated variables.^9^ By contrast, the scanCT displays a more diffuse pattern with predominantly neutral values and fewer large high-correlation blocks (Fig. 2F), suggesting that the features prioritized by scanCT are less redundant and more nearly orthogonal.

We next quantified the distribution of correlation status along the importance ranking. The correlated features of CART and RF are heavily concentrated among the top positions (Fig. 2H, I). Within the top 200, RF and CART include 58 and 64 highly correlated features, respectively. The corresponding importance curves drop steeply, indicating that these clustered predictors receive disproportionately high weights at the expense of unique, low-correlation features. ^19,20^ In contrast, the scanCT curve is noticeably flatter (Fig. 2G), with only two highly correlated features appearing in the top 200 and the majority of top-ranked predictors are low-correlated or uncorrelated, consistent with scanCT favoring features that contribute non-redundant information.

Finally, we evaluated scanCT’s ranking with correlation groups identified by CART and RF. If scanCT captured the collinearity bias of greedy algorithms, features from highly correlated groups in CART and RF would cluster near the top of the scanCT ranking. We oberserved red and orange points are dispersed along the entire scanCT rank axis and intermixed with low-uncorrelated features (blue), with no evidence of systematic accumulation at the top (Fig. 2J, K). Together, these results indicate that scanCT evaluates feature importance based on predictive signal rather than correlation structure, effectively mitigating both cardinality and correlation biases that characterize CART and RF. Results were consistent across B, CD4T, CD8T and T cells (Supplementary Fig. S2, S3, S4 and S5).

**Fig. 3.**
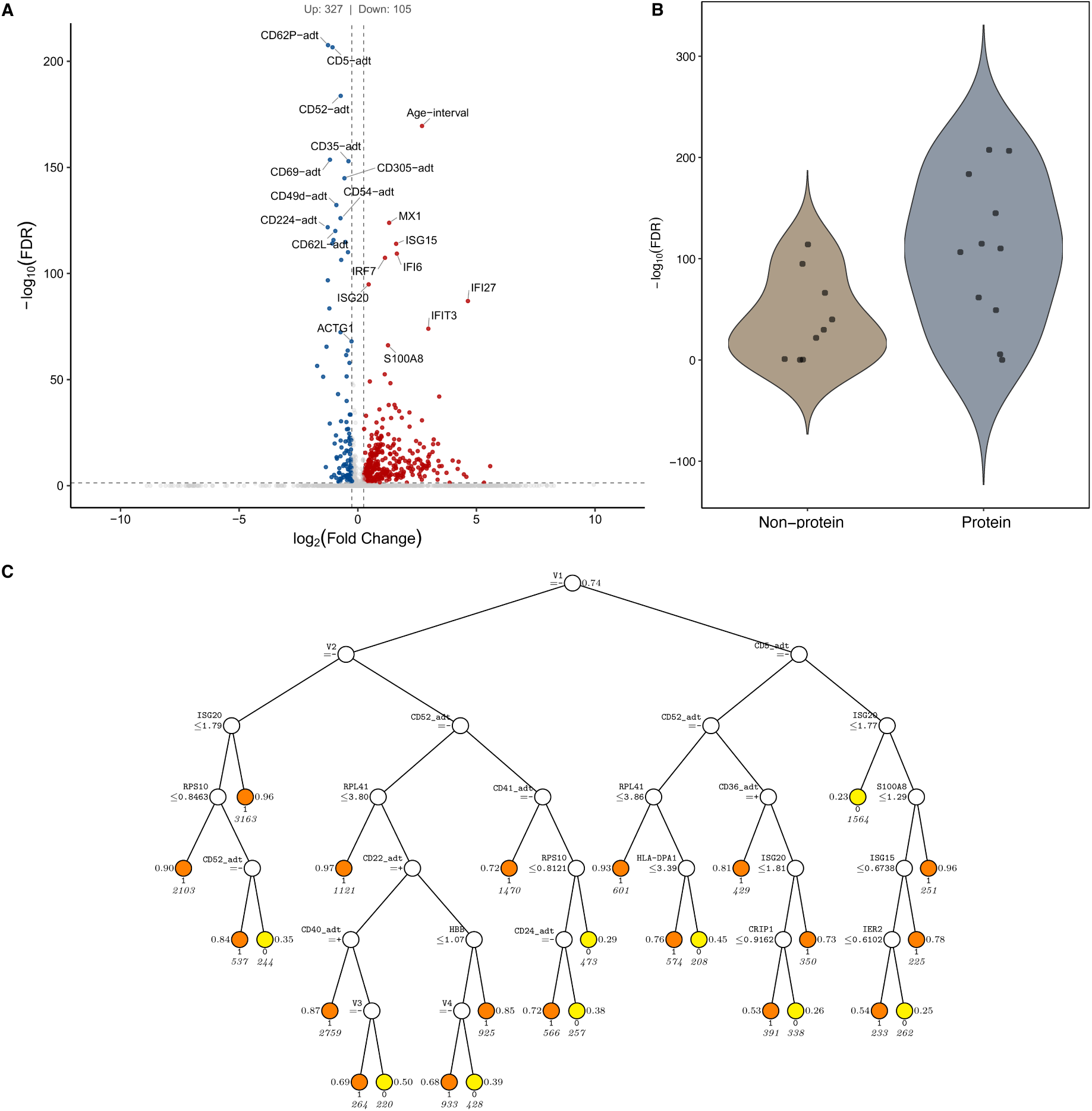
Wilcoxon differential expression and scanCT classification results in B cell Group 2 (B G2). (**A**) Wilcoxon differential expression test comparing groups in B G2; labeled points indicate the top 10 most significant non-protein and protein features. The y-axis represents the negative log-transformed *p*-values (*−* log_10_) derived from the Wilcoxon rank-sum test. (**B**) Wilcoxon significance of the scanCT classification tree splitting variables, summarized by feature type. (**C**) scanCT classification tree for B G2, with feature mapping: V1=CD62P adt, V2=CD268 adt, V3=CD305 adt, V4=CD185 adt. At each split, an observation goes to the left branch if and only if the condition is satisfied. Predicted class and sample size are printed below each terminal node. The proportion of predicted class is printed on the left and right

**Fig. 4.**
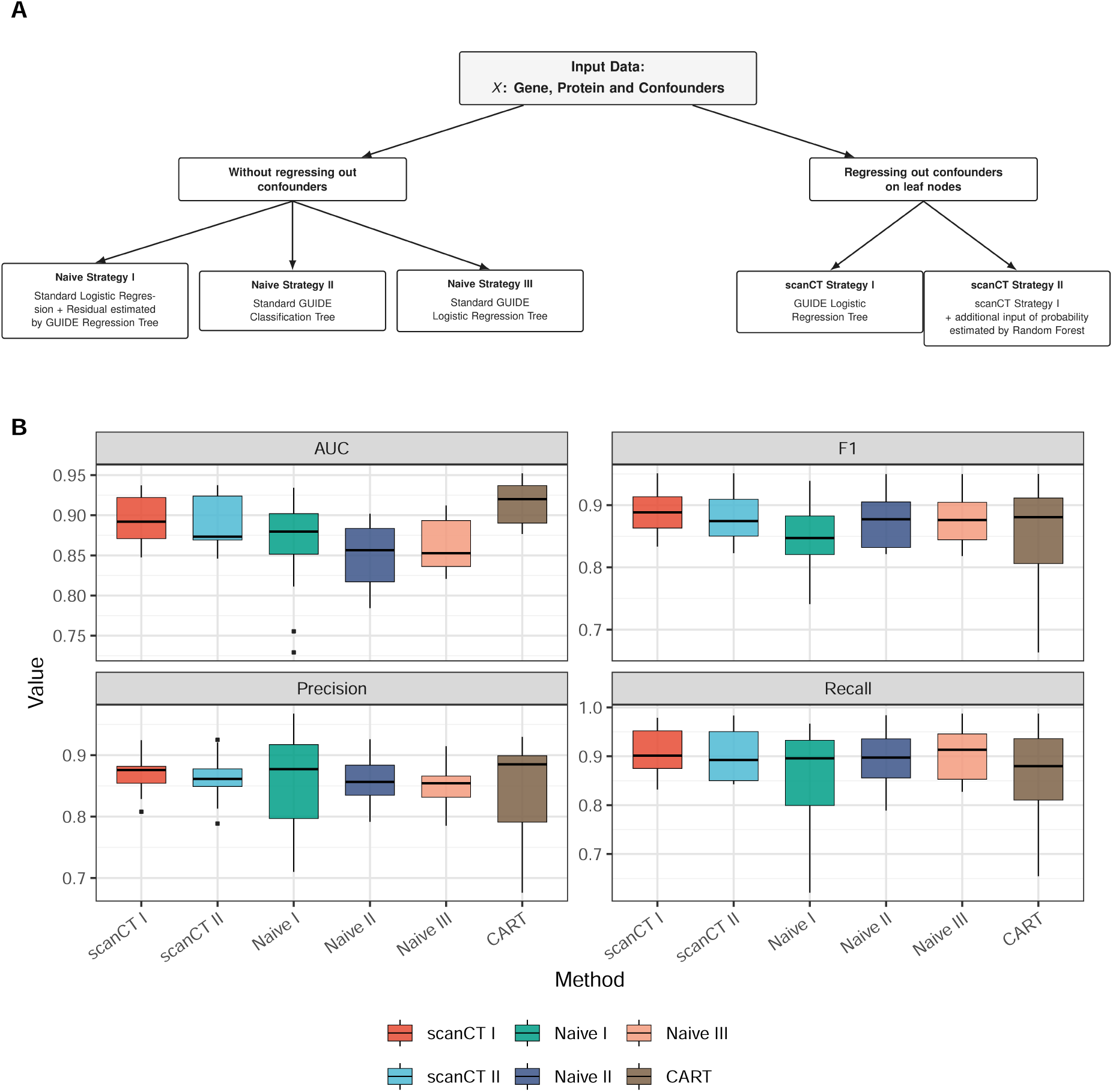
Overview of benchmarking strategies and performance evaluation. (**A**) Schematic workflow of the five modeling strategies categorized by their handling of confounding variables. The framework uses genes, proteins, and confounders as input. The branch without regressing out confounders implements Naive strategies, while the branch regressing out confounders on leaf nodes explicitly adjusts for demographic covariates to isolate disease-specific signals. (**B**) Comparative performance metrics across the five strategies and the standard CART. Bar plots display the Area Under the Curve (AUC), Precision, Recall, and F1 scores. scanCT Strategy I (scanCT I) was selected for final analyses due to its robust performance and superior ability to control for confounders compared to standard CART and Naive implementations.

Collectively, these results indicate that in realistic high-dimensional single-cell settings, conventional tree ensembles can disproportionately allocate variable importance to highly correlated predictors, whereas scanCT distributes importance across a more diverse and low-correlation feature set, which is crucial for downstream biomarker discovery.

### Robust Interpretability and Comparative Benchmarking of scanCT-based Strategies

To elucidate the complex biological mechanisms driving heterogeneity in COVID-19 immune responses, we moved beyond standard differential expression analysis to a structured, interpretable tree approach. While traditional univariate methods effectively identify isolated biomarkers, they often fail to capture the higher-order interactions and conditional dependencies characteristic of cellular signaling pathways. A standard Wilcoxon rank-sum test is commonly used to identify highly significant features. In B-cell Group 2 (B G2), with surface proteins (e.g., CD62P-adt, CD5-adt), the Wilcoxon rank-sum test generally exhibits stronger statistical signals than transcriptomic features (Fig. 3A). However, these visualizations (Volcano and Violin plots) are inherently limited to linear, additive assumptions; they highlight which markers differ but obscure how these markers interact to define cellular states.

In contrast, the scanCT classification tree resolves this limitation by mapping the hierarchical logic of the immune response (Fig. 3C). The tree structure explicitly reveals conditional relationships that are invisible in univariate plots. For instance, the primary stratification is based on CD62P adt (V1), which is identified as the dominant driver of heterogeneity. Importantly, downstream markers such as ISG20 and CD52 adt are selected only within specific branches, suggesting that their biological relevance is context-dependent. This white-box model allows us to disentangle the interplay between surface protein abundance and gene expression, providing a granular view of the B cell dysregulation associated with COVID-19 severity that global differential expression lists cannot offer.

To rigorously validate our modeling approach and ensure that identified biomarkers are disease-specific rather than artifacts of confounding demographic factors, we benchmarked five distinct strategies against the standard CART algorithm. We first categorized these methods into two primary streams: those that model raw data directly (Naive Strategies) and those that rigorously control for confounders such as Age and Sex (scanCT Strategies)(Fig. 4A).

The benchmarking framework takes as input multimodal data and processes them through various regression and tree-construction workflows. Among the evaluated methods, scanCT Strategy I demonstrated superior robustness by consistently achieving high Area Under the Curve (AUC) and F1 scores (Fig. 4B), outperforming standard CART and competing naive implementations.

Crucially, while standard CART is susceptible to variable selection bias that favors continuous variables with more splits, scanCT eliminates this bias through its unbiased curvature tests. Furthermore, by explicitly regressing out Age and Sex at the leaf nodes, scanCT I ensures that classification power is derived from relevant biological signals rather than demographic differences. This makes it the optimal choice for downstream analysis, as it balances high predictive accuracy with the rigorous control of confounding variables necessary for clinical validity.

### Discovery of disease-associated trajectories by scanCT I

#### T-cells

The regression-tree illustrates that T-cell clinical outcomes in the G2 (Healthy vs. Asymptomatic) and G3 (Healthy vs. Severe) cohorts are stratified by distinct but biologically coherent hierarchical checkpoints (Fig. 5).

**Fig. 5.**
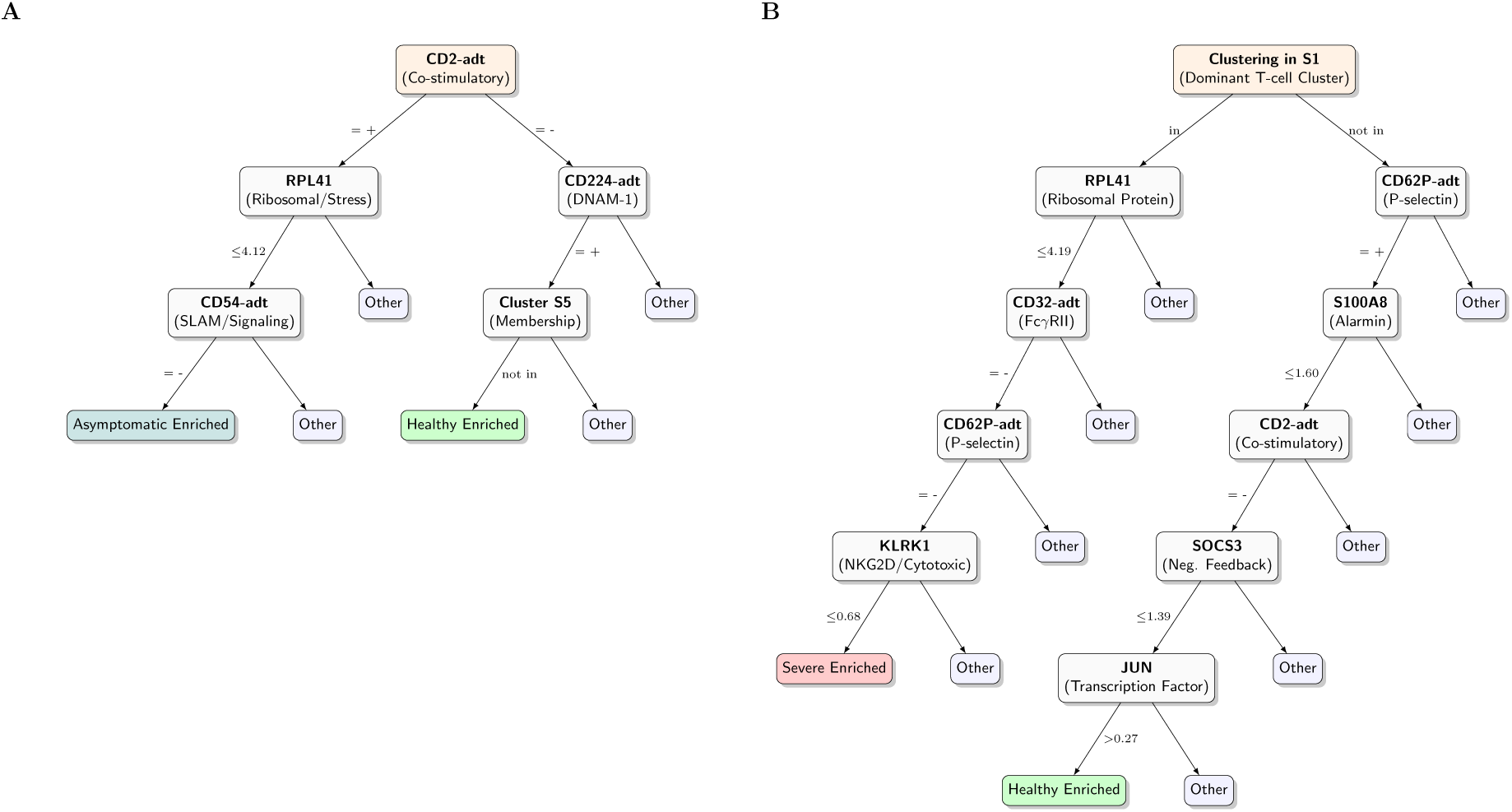
Partial decision-tree structures for T-cell clinical outcome stratification. **(A-B)** scanCT I logistic regression trees for G2 (Healthy vs Asymptomatic) and G3 (Healthy vs Severe). The diagrams visualize the meaningful splits identified by the scanCT algorithm. **Tree Structure:** Orange nodes represent the root and intermediate splitting points. Text within nodes displays the splitting feature (top), its corresponding functional annotation (bottom, in parentheses). **Leaf Nodes:** Terminal nodes are colored according to the dominant clinical outcome (Green: Healthy; Teal: Asymptomatic; Red: Severe). A leaf is labeled as “Enriched” (e.g., Asymptomatic Enriched) only if it contains *≥* 85% of cells from that specific outcome class. **Clustering:** Categorical splits based on cell-type clustering group specific T-cell subpopulations. In G2, cluster *S*_5_ = *{*CD4.CM, CD4.IL22, CD4.Th1, CD8.Naive, CD8.TE, ILC1 3, Treg. In G3, cluster *S*_1_ = *{*CD4.CM, CD4.EM, CD4.Prolif, CD4.Tfh, CD4.Th17, CD8.Prolif, CD8.TE*}*

In the G2 cohort, the primary bifurcation is defined by the co-stimulatory marker CD2 adt (Fig. 5A). This split separates a CD2 adt^+^ compartment which is permissive to infection-associated remodeling, from a CD2 adt^−^ context enriched for baseline homeostatic states. Within the CD2 adt^+^ branch, the trajectory toward an asymptomatic outcome is delineated by a compact rule set anchored in ribosomal function (RPL41) and SLAM-family signaling (CD54 adt). Crucially, the scanCT algorithm identifies coregulated functional modules rather than isolated stochastic markers. The RPL41 split leads a correlated gene module including RPS10, HBB, and CRIP1 (Supplementary Table S2). This co-regulation implies that the asymptomatic State is not defined by the downregulation of a single ribosomal protein, but by a coordinated shift in the cell’s entire translational machinery. This stability offers robust clinical translatability. Identifying any of these correlated high-abundance proteins, or their associated surface markers like CD54 adt (ICAM-1) found downstream, provides alternative candidate biomarkers for patient stratification that are less sensitive to technical dropout than low-abundance transcripts.

In the G3 cohort (Fig. 5B), the tree structure reveals a qualitative shift in the biomarkers associated with severe disease. The severe outcome is predominantly localized within Cluster S1 and is defined by the convergent impairment of biosynthetic capacity and effector sensing. The severe-enriched trajectory follows a path defined by low RPL41, the absence of CD32 adt and CD62P adt, and is finally gated by KLRK1. Here, downregulated RPL41 implies translational constraints consistent with cellular stress, ^21^ while reduced KLRK1 (NKG2D) indicates impaired cytotoxic danger sensing. ^22^ The correlated features for these splits provide a comprehensive list of potential biomarkers that mechanistically validate this trajectory. The KLRK1 split is correlated with a core cytotoxic module including NKG7, CCL5, GZMA, and GNLY (Supplementary Table S2). This explicitly enumerates a cytotoxic collapse signature. Severe disease is not merely the loss of a receptor but the silencing of the entire effector gene program. Furthermore, the correlation of surface CD62P adt (P-selectin) with inflammatory markers highlights a specific failure of extravasation potential, distinguishing these cells from hyper-inflammatory phenotypes.

These trajectories are further contextualized by comparison with baseline models. We generated standard CART decision trees for these same cohorts. In G2, CART similarly identifies CD2 adt as an early splitter. However, the subsequent tree structures degenerate into deep, fragmented chains of single genes, such as IFI6 and FOS, that obscure the underlying cell states (Supplementary Fig. S24). In G3, the distinction is even more profound. The CART algorithm prioritizes Age interval as the primary root split (Supplementary Fig. S25). While Age is a known demographic risk factor, this choice partitions the data based on patient metadata rather than cell-intrinsic biology. In contrast, scanCT prioritizes the cell-state cluster (S1) and mechanistic switches (RPL41, KLRK1). This explicitly isolates the specific T-cell dysfunctions driving pathology regardless of patient Age. This confirms that scanCT isolates a parsimonious set of interpretable, biologically actionable switches that map directly to the immunobiology of COVID-19.

#### CD4T-cells

CD4T-cell outcomes in G2 and G3 are separated by compact, interpretable checkpoints that map onto expected COVID-19 immunobiology (Fig. 6).

**Fig. 6.**
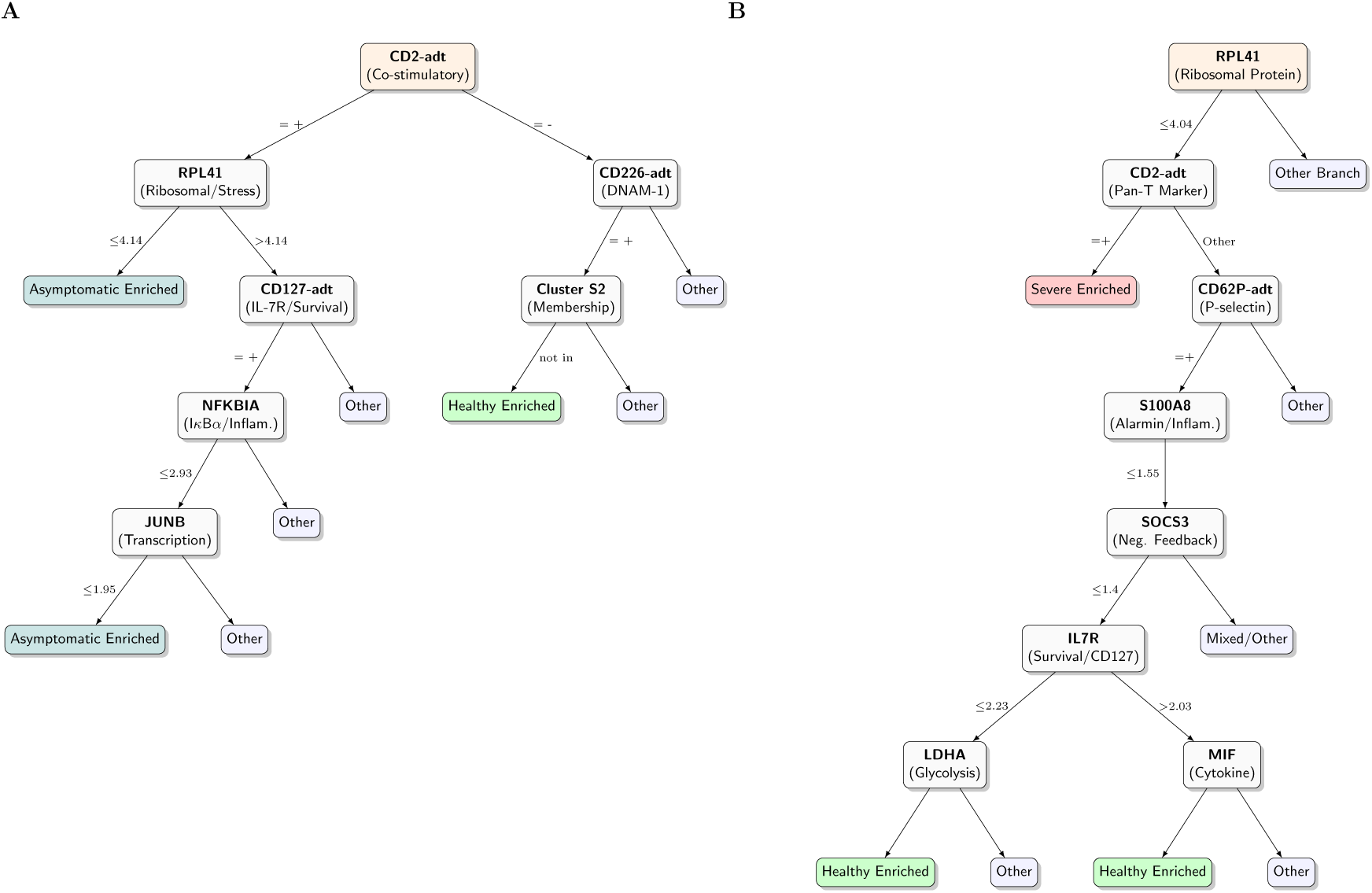
Partial decision-tree structures for CD4T-cell clinical outcome stratification. **(A-B)** scanCT I logistic regression trees for G2 (Healthy vs Asymptomatic) and G3 (Healthy vs Severe). The diagrams visualize the meaningful splits identified by the scanCT algorithm. **Tree Structure:** Orange nodes represent the root and intermediate splitting points. Text within nodes displays the splitting feature (top) and its corresponding functional annotation (bottom, in parentheses). **Leaf Nodes:** Terminal nodes are colored according to the dominant clinical outcome (Green: Healthy; Teal: Asymptomatic; Red: Severe). A leaf is labeled as ”Enriched” only if it contains *≥* 85% of cells from that specific outcome class. **Clustering:** Categorical splits based on cell-type clustering group-specific CD4^+^ T-cell subpopulations. In G2, cluster *S*_2_ = *{*CD4.CM, CD4.EM, CD4.Prolif, CD4.Tfh*}*.

In the G2 cohort (Fig. 6A), the initial split on CD2 adt separates a CD2^+^ co-stimulatory compartment permissive to infection-associated remodeling from a CD2^−^ context enriched for baseline states. Within the CD2^+^ branch, the dominant asymptomatic trajectory is captured by a ribosomal-stress gate. scanCT identifies RPL41 as the primary switch. Crucially, this feature proxies for a broad co-regulated stress module explicitly enumerated in the node, including HBB (hemoglobin scavenging/oxidative stress), CRIP1, SAT1, and the interferon-stimulated gene ISG15 (Supplementary Table S3). This provides stable, multigene evidence that mild infection associates with a stress-conditioned program rather than a single marker loss. A second asymptomatic route arises when cells maintain translational capacity (RPL41^high^) but are stratified by survival and feedback regulation (CD127 adt, NFKBIA, JUNB). This trajectory identifies a specific cell state characterized by controlled activation shaped by inflammatory negative-feedback circuitry. By contrast, healthy enrichment is localized to the alternative branch marked by DNAM-1 (CD226 adt) and is excluded from Cluster S2. This suggests that healthy samples preferentially occupy a stable homeostatic cluster rather than infection-remodeled compartments.

In the G3 cohort(Fig. 6), the hierarchy shifts toward a distinct architecture in which the Severe outcome is isolated early via an RPL41-rooted branch, coupled with reduced CD2 adt. The correlated features for CD2 adt reveal a striking collapse of the AP-1 transcription factor complex (FOS, JUN, JUNB) and factors maintaining quiescence and identity (KLF2) (Supplementary Table S3). This implies that severe disease in CD4T cells involves a double hit: translational constraint (RPL41^low^) followed by the loss of core transcriptional identity. A regulated inflammatory–metabolic module instead organizes the healthy recovery trajectory. The split on S100A8 proxies for a complete alarmin cassette (S100A9, LYZ), while the downstream SOCS3 split correlates with metabolic regulators like LDHA. This indicates that recovery-compatible states retain coordinated metabolic flexibility and feedback control, distinguishing them from the effector collapse seen in severe disease.

These mechanistic trajectories are less transparent in baseline models. The standard CART decision trees struggle to isolate cell-intrinsic drivers from patient demographics. In the G2 cohort (Supplementary Fig. S18), CART prioritizes Patient Sex as the root split before accessing biological markers. Similarly, in G3 (Supplementary Fig. S19), while CART identifies the ribosomal signal, it relies heavily on Age interval and broad interactions with Sex to partition the data. This demographic dominance obscures the universal cellular checkpoints. In contrast, the scanCT I trees isolate a parsimonious set of interpretable biological and define clinically meaningful trajectories applicable across demographic groups by switching co-stimulation (CD2), translational stress (RPL41/HBB), and metabolic feedback (S100A8, SOCS3).

#### CD8T-cells

CD8T-cell outcomes in G2 and G3 are governed by distinctive hierarchical checkpoints that map to coherent functional modules rather than isolated marker fluctuations (Fig. 7).

**Fig. 7.**
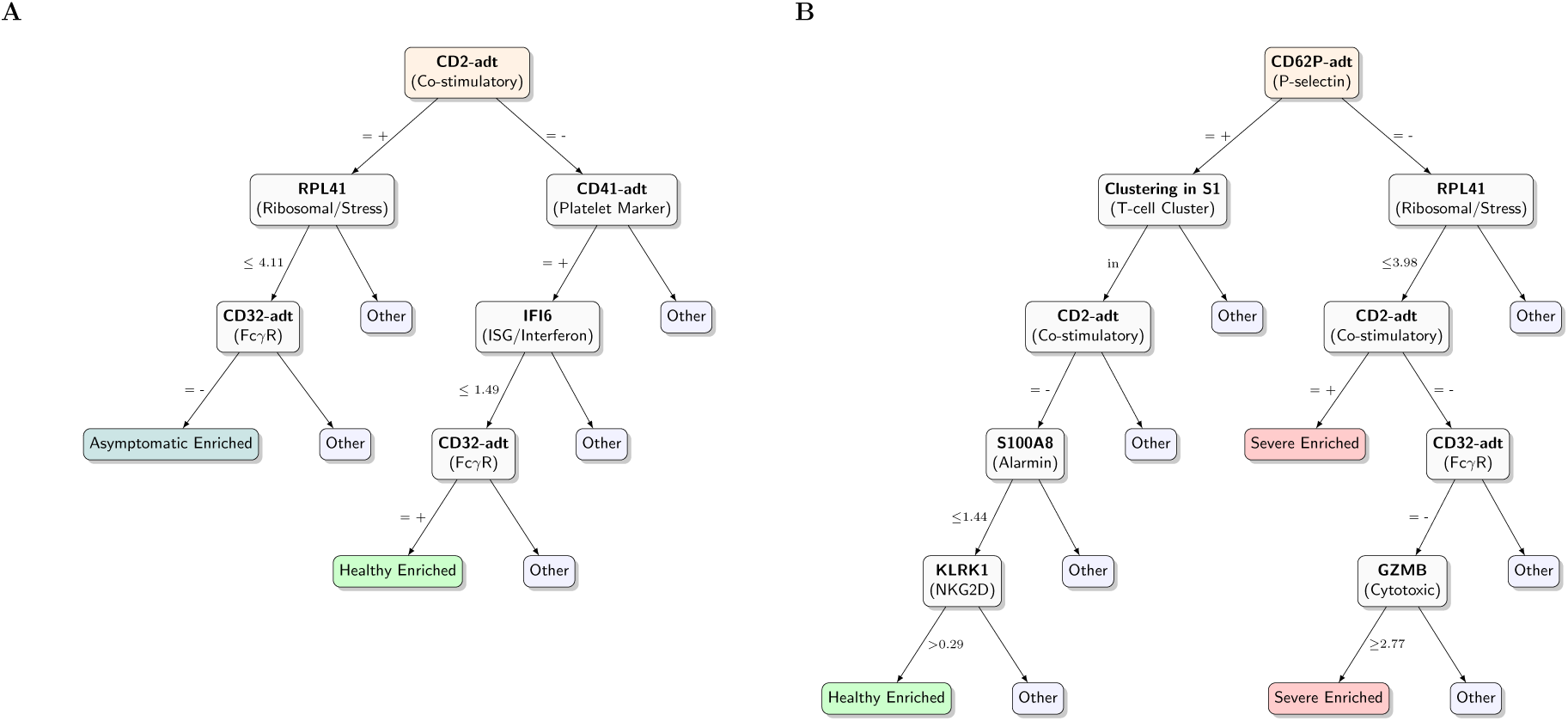
Partial decision-tree structures for CD8T-cell clinical outcome stratification. **(A-B)** scanCT I trees for G2 (Healthy vs Asymptomatic) and G3 (Healthy vs Severe). The diagrams visualize the top-ranking splits identified by the scanCT algorithm. **Tree Structure:** Orange nodes represent the root and intermediate splitting points. Text within nodes displays the splitting feature (top) and its corresponding functional annotation (bottom, in parentheses). **Leaf Nodes:** Terminal nodes are colored according to the dominant clinical outcome (Green: Healthy; Teal: Asymptomatic; Red: Severe). A leaf is labeled as ”Enriched” only if it contains *≥* 85% of cells from that specific outcome class. **Clustering:** Categorical splits based on cell-type clustering group-specific CD8^+^ T-cell subpopulations. In G3, cluster *S*_1_ = *{*CD8.EM, CD8.Naive*}*.

In the G2 cohort (Fig. 7A), the dominant bifurcation occurs at CD2 adt. This separates a CD2 adt^+^ co-stimulatory compartment, permissive to infection-associated remodeling, from an alternative branch characterized by platelet-marker context (CD41 adt) and interferon-stimulated signaling (IFI6). Within the CD2 adt^+^ branch, the specific trajectory to asymptomatic enrichment is gated by RPL41 and CD32 adt. Crucially, the RPL41 split acts as a proxy for a stabilized stress-effector module and it is tightly co-regulated with both stress markers (FOS, HBB) and key cytotoxic effectors (GZMB, PRF1, NKG7) (Supplementary Table S4). This indicates that asymptomatic infection promotes a controlled state where translational constraints (RPL41) are biologically coupled to effector readiness, preventing the runaway inflammation seen in severe disease.

In the G3 cohort (Fig. 7B), the hierarchy shifts toward a failure architecture rooted in an interaction context (CD62P adt) and translational stress (RPL41). Severe outcome is concentrated along the CD62P adt^−^ and RPL41 branch and is specified early by reduced co-stimulatory context (CD2 adt). The final refinement for severity involves CD32 adt and GZMB (threshold 2.77). Here, the GZMB split is synonymous with a broad cytotoxic collapse and it correlates with the loss of GNLY, PRF1, NKG7, and GZMH (Supplementary Table S4). This explicitly identifies the severe State as a double-negative dysfunction. Cells fail to maintain biosynthetic integrity (low RPL41) and simultaneously lose their core cytotoxic machinery (low GZMB module), rendering them immunologically inert. ^22^ Conversely, the healthy-enriched trajectory is preserved within the S1 cluster. It is defined by the retention of cytotoxic sensing (KLRK1/NKG2D) and the absence of alarmins (S100A8), confirming that recovery requires maintained surveillance capacity.

These mechanistic trajectories are further clarified by comparison to baseline models. Standard CART trees of G2 produce a fragmented structure (Supplementary Fig. S20), splitting first on CD2 adt but diluting predictive power across stochastic downstream markers like IFI6 and technical artifacts. Most strikingly, the CART tree of G3 (Supplementary Fig. S21) selects Age interval as the root splitting variable. While Age is a potent demographic covariate, this splits the data based on patient risk rather than T-cell intrinsic biology. By contrast, scanCT successfully isolates CD62P adt and RPL41 as the primary determinants, distinguishing the specific cellular mechanisms of progression (adhesion and translational failure) independent of patient Age. This confirms scanCT’s utility for discovering translatable biomarkers, such as the KLRK1 cytotoxic-sensing axis, that offer specific therapeutic targets beyond broad demographic risk stratification.

#### Mononuclear-cells

Mononuclear-cell clinical outcomes are organized by coherent hierarchical checkpoints that align with established COVID-19 immunobiology, specifically transitioning from interferon-regulated states in mild infection to inflammatory dysregulation in severe disease (Fig. 8).

**Fig. 8.**
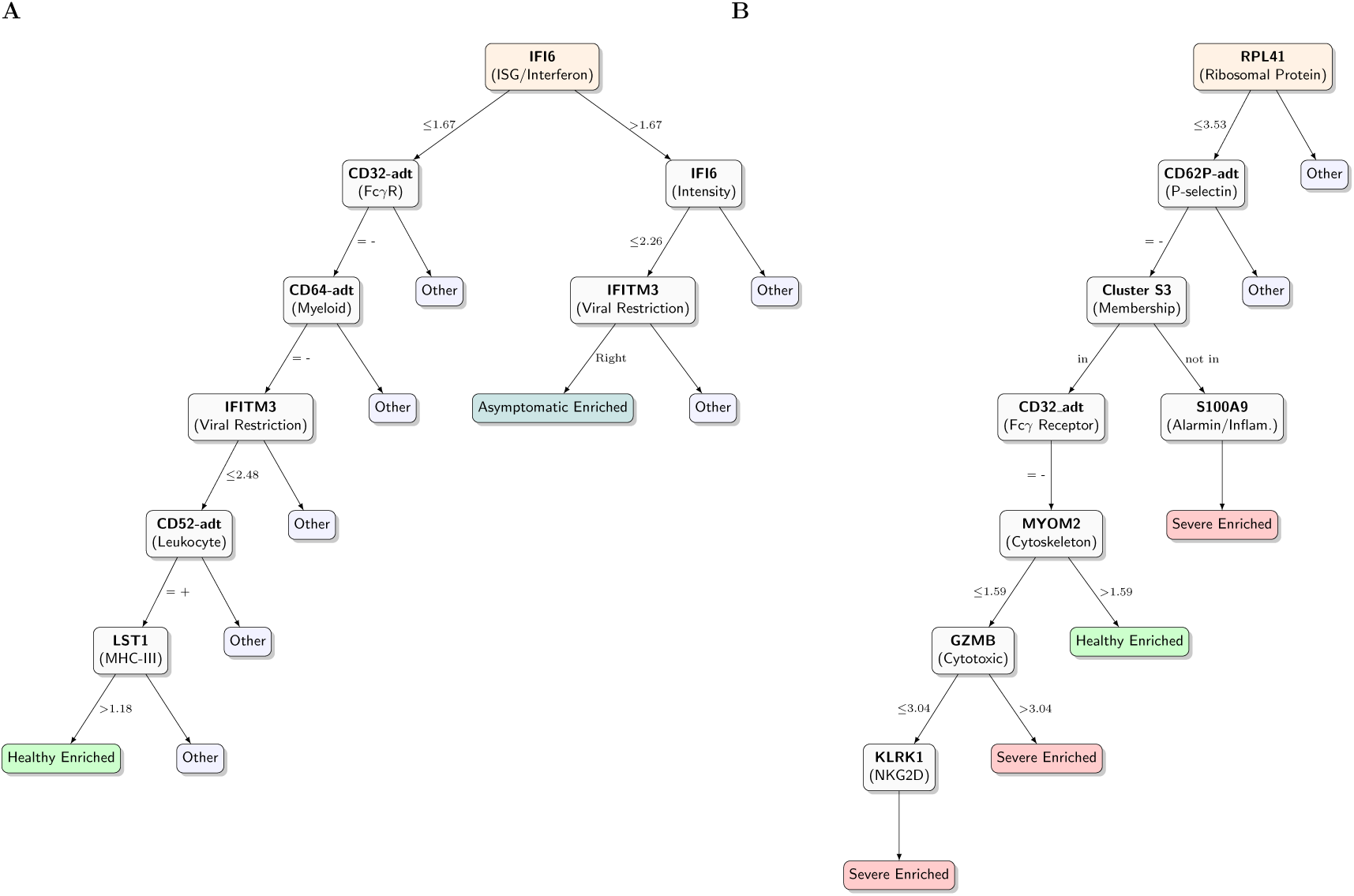
Partial decision-tree structures for Mononuclear cell clinical outcome stratification. **(A-B)** scanCT I trees for G2 (Healthy vs Asymptomatic) and G3 (Healthy vs Severe). The diagrams visualize the top-ranking splits identified by the scanCT algorithm. **Tree Structure:** Orange nodes represent the root and intermediate splitting points. Text within nodes displays the splitting feature (top) and its corresponding functional annotation (bottom, in parentheses). **Leaf Nodes:** Terminal nodes are colored according to the dominant clinical outcome (Green: Healthy; Teal: Asymptomatic; Red: Severe). A leaf is labeled as ”Enriched” only if it contains *≥* 85% of cells from that specific outcome class. **Clustering:** Categorical splits based on cell-type clustering group specific mononuclear subpopulations. In G3, cluster *S*_3_ = *{*CD14 Mono, CD16 Mono, cDC, pDC, Platelet, NK*}*.

In the G2 cohort (Fig. 8A), the primary bifurcation occurs at IFI6, establishing interferon-stimulated signaling as the dominant axis of separation. The healthy-enriched trajectory is embedded in a myeloid/-monocyte context specified by low CD32 adt (Fc*γ*RII), the absence of activated markers CD64 adt and IFITM3, and the presence of LST1. Notably, LST1 acts as a stable anchor for a homeostatic myeloid module correlated with SERPINA1, FTH1, and CD68 (Supplementary Table S5). This module confirms that the healthy State maintains a quiescent, structured myeloid compartment rather than the infection-remodeled, interferon-high state seen in the alternative branch. Conversely, asymptomatic enrichment follows the IFI6-high trajectory. It is further gated by IFITM3, suggesting that mild infection is captured as a controlled antiviral program where the intensity of viral restriction (IFITM3) calibrates the clinical outcome.

In the G3 cohort (Fig. 8B), the tree emphasizes a distinct architecture centered on translational stress and inflammatory remodeling. Severe outcome is concentrated along an RPL41-rooted branch coupled to a CD62P adt context. This outcome is further stratified by cluster membership (Cluster S3) and Fc*γ*R-associated context (CD32 adt). The trajectory culminates in a cytotoxic-program gate defined by GZMB and KLRK1. Here, the low GZMB branch explicitly enumerates a cytotoxic collapse. It correlates with the loss of GNLY, NKG7, PRF1, and GZMA (Supplementary Table S5), identifying a severe-specific state where NK/effector cells lose their lytic machinery while retaining the stress phenotype. A secondary severe-enriched route is defined by the alarmin axis (S100A9), which correlates with CD14, VCAN, and FCN1 (Supplementary Table S5). This cleanly captures the emergency myelopoiesis signature, distinguishing the dysfunctional, inflammatory monocytes in severe COVID-19.^23^

These mechanistic trajectories are obscured in baseline models. While the CART tree of G2 (Supplementary Fig. S22) correctly identifies the early IFI6 split, it rapidly fragments into complex chains of Age interval and stochastic technical splits. Most critically, the CART tree for G3 (Supplementary Fig. S23) also selects Age interval as the root splitting variable. This demographic prioritization masks the cellular mechanism. Essentially, CART sorts patients by age risk, whereas scanCT sorts cells by functional State. By prioritizing cell-intrinsic switches like translational integrity (RPL41), alarmin activation (S100A9), and cytotoxic capacity (GZMB), scanCT reveals the specific immunologic failures driving severe disease. This offers a direct map to potential therapeutic targets that is independent of patient demographics.

#### B-cells

The scanCT I trees for B cells define compact, mechanistically interpretable checkpoints that separate mild infection-associated remodeling from severe disease states through interactions with platelets, interferon signaling, and B-cell survival or antigen-presentation modules (Fig. 9).

**Fig. 9.**
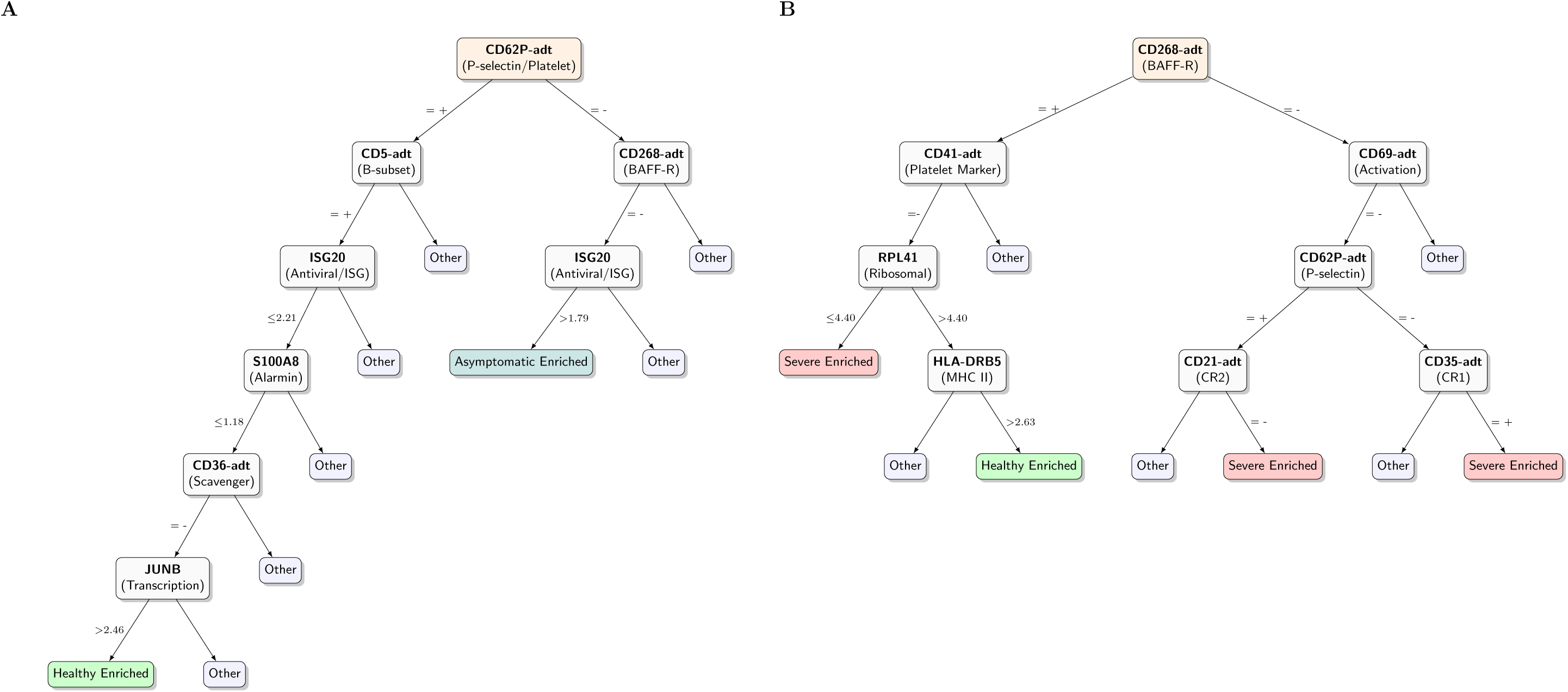
Partial decision-tree structures for B-cell clinical outcome stratification. **(A-B)** scanCT I trees for G2 (Healthy vs Asymptomatic) and G3 (Healthy vs Severe). The diagrams visualize the top-ranking splits identified by the scanCT algorithm. **Tree Structure:** Orange nodes represent the root and intermediate splitting points. Text within nodes displays the splitting feature (top) and its corresponding functional annotation (bottom, in parentheses). **Leaf Nodes:** Terminal nodes are colored according to the dominant clinical outcome (Green: Healthy; Teal: Asymptomatic; Red: Severe). A leaf is labeled as ”Enriched” only if it contains *≥* 85% of cells from that specific outcome class.

In the G2 cohort (Fig. 9A), the top split at CD62P adt suggests that B-cell outcome stratification is strongly conditioned on an adhesion and platelet-associated interaction milieu. Within the CD62P adtpositive context, the healthy-enriched State is defined by a CD5 adt subset and ISG20 antiviral checkpoint. It is further characterized by low levels of the inflammatory marker S100A8 and scavenger receptor CD36 adt. This profile is consistent with a regulated activation background rather than an infection-driven inflammatory program. By contrast, asymptomatic enrichment aligns with a CD62P adt-negative context structured by CD268 adt (BAFF-R) and limited ISG20 engagement. This pattern suggests that clinically mild infection is captured by a BAFF-R–conditioned survival state with minimal engagement of the acute antiviral ISG module.

In the G3 cohort (Fig. 9B), outcome separation is anchored by CD268 adt (BAFF-R), highlighting B-cell survival or tonic signaling as a key context for severe disease trajectories. ^24^ Severe enrichment is observed in a platelet-associated context (CD41 adt) characterized by low RPL41. ^25^ This is consistent with stress-associated translational repression and systemic inflammatory burden. A second severe-associated mode is defined by an activation/complement-receptor architecture involving CD69 adt and complement receptors CD21 adt and CD35 adt, which is consistent with altered immune-complex handling and complement-linked B-cell remodeling described in severe infection. Conversely, healthy enrichment corresponds to a high HLA-DRB5 (MHC II) state within the platelet-associated BAFF-R branch. This indicates preserved antigen-presentation capacity and immune coordination compatible with the healthy attractor. Comparison to baseline models emphasizes the utility of scanCT trees for mechanistic interpretation, even when root splits align. In G3 (Supplementary Fig. S17), the CART tree also identifies CD268 adt as the root split, validating BAFF-R as the primary discriminator. However, the downstream CART structure fragments into redundant gene splits, such as RPS10 and ZFP36, without clearly delineating the functional states. scanCT, by contrast, explicitly resolves the severe trajectory into a translational stress arm (low RPL41) and a complement activation arm (CD21 adt/CD35 adt). This provides a structured biological map of the specific dysregulation modes, including coagulopathic interactions and inflammatory exhaustion, that define severe B-cell pathology.

### Micro-level node analysis of confounder adjustment: a case study of CD8T cells

To illustrate the local behavior of the scanCT logistic regression tree, we performed a node-level analysis for the CD8T-cell subset in G2.

Terminal node 182 captures a distinct genomic subpopulation defined by specific protein and gene expression signatures (Fig. 10). This node serves as a representative example from the structure shown in Fig. S10. The node-specific logistic regression coefficients for Age and Sex, together with the corresponding *t*-test *P* values across terminal nodes, provide a direct measure of how reliably each node supports inference on confounder effects.

**Fig. 10.**
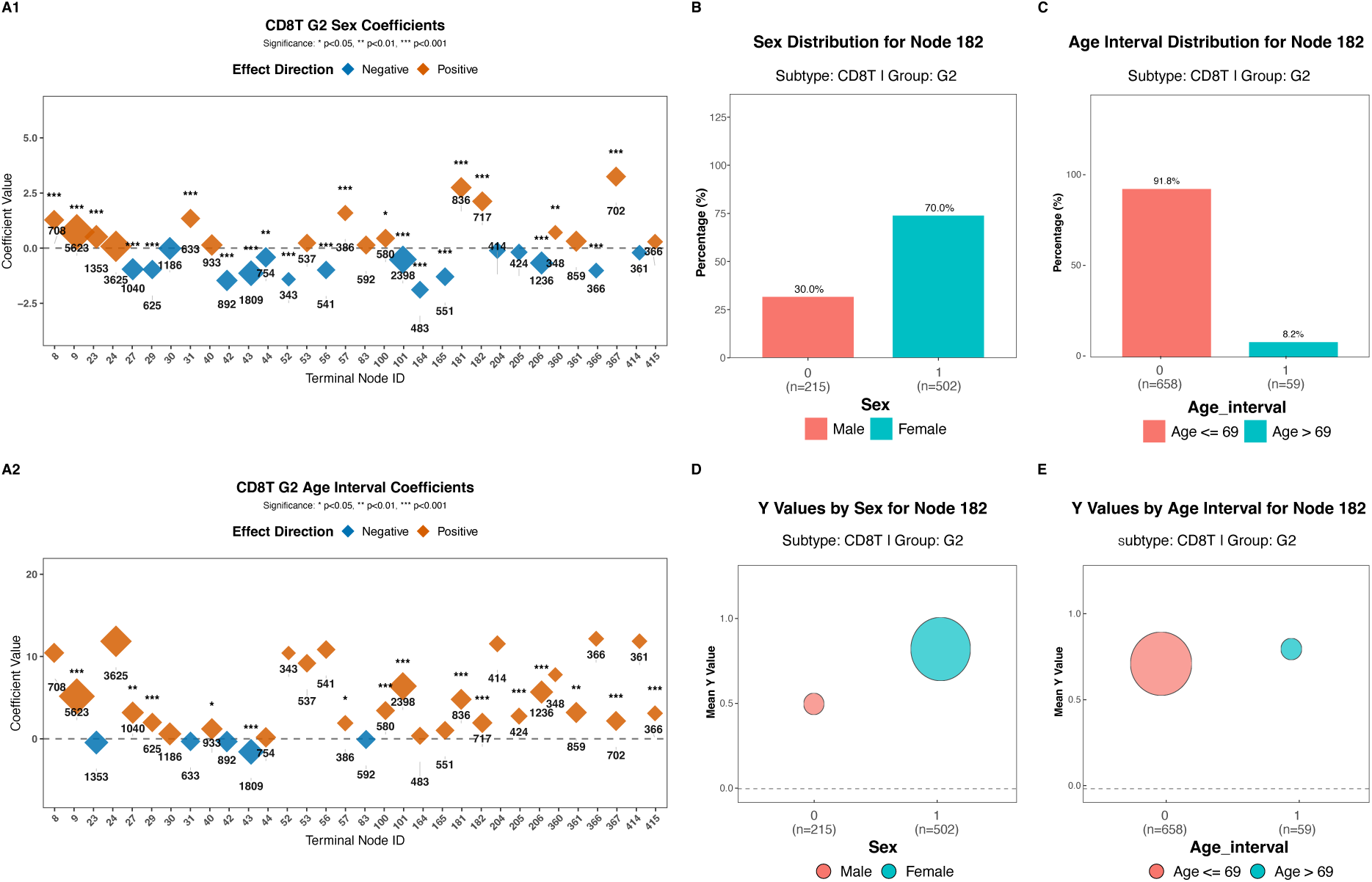
Micro-level analysis of CD8T subset (Group 2, Node 182). **(A)** Terminal node coefficients for Age and Sex across all nodes in the CD8T/G2 tree, showing the variability of confounder effects. **(B, C)** Distribution of Sex and Age, showing a skew towards females and younger patients. **(D, E)** Mean severity scores (Y) stratified by Sex and Age, highlighting higher severity in females and older individuals within this specific subset.

The node is strongly skewed toward female patients (70.0% female) and younger individuals (≤ 69 years old, 91.8%) (Fig. 10B, C). Within this specific molecular context, demographic traits exert pronounced and synergistic effects on severity. Older patients (*>* 69 years old) exhibit a higher mean severity score (0.83) compared to younger patients (0.74) (Fig. 10E). Similarly, females exhibit markedly higher severity (mean 0.85) compared to males (mean 0.52) (Fig. 10D). This indicates a synergistic relationship where both risk factors, female sex and older Age, align to drive higher disease severity within this subpopulation.

Critically, these local dynamics underscore the limitations of global adjustment methods (Naive I– III). A global regression would average these effects across the entire dataset. This could dilute the strong sex-dependency observed here or apply it inappropriately to other nodes where the effect is absent or reversed. By fitting node-specific models, scanCT explicitly unmasks local confounder variations. This ensures that adjustment is tailored to each cell subset’s biological context.

## Discussion

Here, we evaluated scanCT as an interpretable framework for modeling high-dimensional single-cell multi-omics data in the presence of heterogeneous feature types and demographic confounding. Across a cohort-scale dataset with mixed continuous gene-expression and binarized ADT protein features, we found that scanCT yields feature rankings and decision rules that are substantially less distorted by common artifacts of tree-based learning. Collectively, our results position scanCT as a practical alternative to conventional tree ensembles when robust biomarker prioritization, transparent decision logic, and calibrated risk estimation are required.

A central finding is that scanCT mitigates feature selection bias driven by predictor cardinality. Under a permutation null, CART and Random Forest assigned disproportionately high importance to high-cardinality gene-expression features. This indicates that apparent ”top markers” can emerge even when no biological signal is present. In contrast, scanCT’s importance scores were effectively decoupled from the number of unique values. This supports its structural neutrality in split selection. This property is particularly consequential in multi-omics settings, where modalities naturally differ in sparsity, dynamic range, and discretization. Without bias control, downstream biological interpretation may conflate statistical granularity with mechanistic relevance.

We further show that scanCT reduces correlation-driven redundancy among top-ranked features. Whereas CART and Random Forest preferentially returned tightly collinear groups, consistent with the known inflation of importance for correlated predictors, scanCT produced a more diverse set of high-ranking variables with fewer dense correlation blocks. In practice, this implies that scanCT can prioritize a feature set that is less dominated by redundant proxies. This is advantageous for biomarker discovery workflows where experimental validation resources are limited and non-overlapping candidates are preferred.

Despite its emphasis on interpretability, scanCT achieved competitive predictive performance relative to CART. While CART typically delivered some higher raw discrimination metrics, scanCT single-tree models retained much of the predictive signal while providing explicit decision rules that map molecular measurements to clinically meaningful severity strata. Importantly, this transparency supports hypothesis generation. The tree topology highlights a small number of splits and thresholds that define coherent subgroups. This enables direct biological interrogation that is difficult to extract from black-box ensembles.

A key contribution of this work is the use of scanCT logistic regression trees to adjust for demographic confounders while preserving interpretability. By fitting node-specific logistic models with Age and Sex as covariates within molecularly defined subpopulations, scanCT produced better-calibrated probability estimates than naive adjustment approaches. This suggests that confounding effects are not uniform across the feature space. Global adjustments can dilute strong local effects by averaging across heterogeneous subgroups. In contrast, scanCT’s hybrid structure allows molecular features to define clinically and molecularly coherent partitions while quantifying how confounders affect the outcome.

Several extensions would further strengthen the scope and generality of our framework. First, while we demonstrate confounder adjustment for two demographic variables (Age and Sex), future work should incorporate continuous and multi-category confounders, such as treatment and batch effects. It should also systematically assess sensitivity to alternative encodings and threshold choices, including ADT binarization schemes and age cut-points. Second, our evaluation targets specific correlation and cardinality regimes. Broader stress testing across additional datasets, modalities, and preprocessing pipelines (normalization, integration strategies, and missingness patterns) will be necessary. This testing will delineate the conditions under which scanCT provides the most significant advantage and those in which ensemble methods remain preferable. Third, extending scanCT logistic regression trees to directly model ordinal or multi-class outcomes would better reflect clinical severity scales. This would also reduce information loss from binarization. This direction includes developing and benchmarking ordinal-logistic node models and establishing appropriate calibration diagnostics for multi-class risk estimation.

## Methods

### Dataset and experimental design

We analyzed a high-dimensional single-cell multi-omics dataset comprising *N* = 110 individuals with COVID-19 or a healthy status. For each subject, approximately 5,000 peripheral blood mononuclear cells were profiled, yielding 2,587 features per cell, including continuous gene expression and binarized protein abundances measured by antibody-derived tags (ADT).

The primary clinical outcome is COVID-19 severity, encoded on an ordinal scale *Y* 0, 1, 2, 3, 4, 5 corresponding to Healthy, Asymptomatic, Mild, Moderate, Severe, and Critical, respectively:

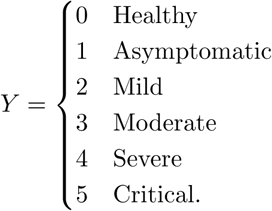

To probe model behaviour across different disease contrasts and signal strengths, we defined four comparison scenarios (G1–G4; Table 1), ranging from broad case–control discrimination (Healthy vs any COVID-19) to fine-grained separation of severe/critical cases.

Cell types were defined by aggregating the original fine-grained clusters into major immune compartments using the study-provided annotations and canonical marker genes (Supplementary Table S1). The Mononuclear Phagocytes and HSPC compartment included dendritic cell subsets, monocytes, platelets, hematopoietic stem and progenitor cell clusters, and natural killer (NK) cell clusters. The T-cell compartment comprised CD4T and CD8T subsets, additional T-lineages (NKT, Treg, *γδ*T and MAIT), and innate lymphoid cells (ILCs). B-lineage cells were grouped into B-cell and plasma-cell clusters. For analyses requiring finer resolution, we also considered CD4T and CD8T cells as dedicated compartments. All predictive models were fitted separately within each immune compartment and comparison group to capture lineage-specific molecular associations with clinical severity.

### GUIDE classification tree

GUIDE classification trees build binary decision trees by recursively partitioning the data while avoiding the variable selection bias of impurity-based methods. ^13^ At each node, a contingency table chi-squared test is used to screen candidate splitting variables. For each predictor *X*, we construct a contingency table with *X* in the columns and the categorical response *Y* in the rows. Ordinal predictors are first grouped into 3–4 intervals at sample quantiles, and categorical predictors use their observed categories; missing values are treated as an additional category.

The splitting variable is chosen as the *X* with the smallest *p*-value from the chi-squared test, which is approximately independent of the number of possible split points and therefore reduces selection bias toward high-cardinality variables. Conditional tests are used to detect local interactions when appropriate. Once the splitting variable is selected, a univariate search over all admissible splits of the form *X ≤ c* (ordinal) or *X ∈ A* (categorical) is performed to minimize the sum of squared deviations of the node-wise class indicator vectors around their means. For ordinal *X*, the search considers splits such as *X c* and *X > c* , with *c* ranging over midpoints of ordered values and with missing values routed optimally to one branch. For categorical *X*, the search is over all subsets *A* of categories (including the missing-value category).

Trees are grown recursively until a large maximal tree is obtained, subject to a minimum node size, and then pruned using cost-complexity pruning to maximise a 10-fold cross-validation estimate of classification accuracy. Missing values are never imputed: at each split, GUIDE determines whether observations with missing *X* are sent to the left or right child to optimise node fit. Variable importance for bare classification trees is defined as the total chi-squared association of each predictor with the response over all split-selection tests up to a specified interaction order (here, 4th order), as described below.

### GUIDE logistic regression tree

GUIDE logistic regression trees (LRTs) extend classification trees by fitting an ordinary logistic regression (OLR) model in each terminal node while retaining GUIDE’s unbiased split-selection mechanism. ^26^ The goal is to allow genomic features *X* to determine the data partitioning, while confounders *Z* serve as covariates in node-specific logistic models.

The algorithm proceeds in several steps. First, a GUIDE forest (an ensemble of least-squares regression trees) is fitted to obtain a preliminary probability estimate *p̃*(*x*) for each observation. Within a given node, missing predictor values are temporarily imputed by node-wise means, and an OLR model is fitted to yield a local estimate *p*^(*x*). Pseudo-residuals *r* = *p̃*(*x*) *p*^(*x*) are then computed and converted to a binary sign indicator, which serves as a surrogate response for split selection.

For each predictor *X*, a contingency table is formed between the discretised *X* (with missing coded as a separate category) and the sign of *r*, and a chi-squared test is performed. The predictor with the most significant test is chosen to split the node. Given the selected *X*, GUIDE searches over candidate thresholds *c* (ordinal) or subsets of categories *A* (categorical) and chooses the split that minimizes the sum of residual deviances of the OLR models fitted in the resulting child nodes. Mean imputation is used only during model fitting; for splitting, observations with missing *X* are explicitly sent to the child that minimizes total deviance. The tree is grown recursively and pruned using cost-complexity pruning to minimize a cross-validated estimate of deviance. In our confounder-adjustment analyses, *X* comprised genomic features used for splitting, whereas confounders *Z* (Age, sex) were included as covariates in all node-specific logistic models.

### GUIDE importance scores

We used the GUIDE variable importance procedure, which combines chi-squared association tests with permutation-based standardisation to yield scores that are approximately unbiased across variable types. ^27^ For each intermediate node *t* with sample size *n_t_*, a constant is fitted to the response, and residuals are recoded into a binary class variable *Z_t_* (e.g., above vs below the node mean). Continuous and ordinal predictors *X_k_* are discretised into categorical variables *X*^′^ (with missing values treated as an explicit category), and for each pair (*X*^′^ *, Z_t_*) a contingency table chi-squared test is performed.

Let *p*_1_(*k, t*) be the *p*-value for predictor *X_k_* at node *t*. GUIDE constructs an unadjusted importance contribution

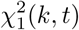

as the (1 − *p*_1_(*k, t*))-quantile of the chi-squared distribution with 1 degree of freedom, and defines

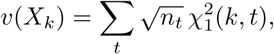

Where the sum is over all intermediate nodes (with additional steps to account for local pairwise interactions when present), to correct for the tendency of some variable types to generate larger test statistics under the null, GUIDE estimates the expected unadjusted score *v*(*X_k_*) by repeatedly permuting the response *Y* while holding the predictors fixed. Specifically,

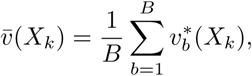

where *v*^∗^(*X_k_*) is computed from the *b*-th permutation. The final standardised importance score is

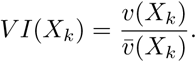

In our experiments, we used this standardised score to rank features, reporting the mean and variability of *V I*(*X_k_*) across 20 permutation replicates.

### Tree strategies and evaluations

We evaluated five strategies for adjusting for demographic confounders *Z* (Age and Sex), including classification and logistic models, ranging from naive baselines to GUIDE-based approaches. Throughout, *X* denotes genomic and protein predictors and *Y* ∈ {0, 1} the binary outcome.

#### Naive Strategy I: global residual adjustment

A global logistic regression model is first fitted for *Y* on *Z*,

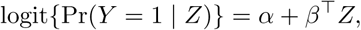

yielding baseline probabilities *p*^*_Z_* = Pr(*Y* = 1 | *Z*). We then compute residuals *Y*_res_ = *Y* − *p*^*_Z_* and fit a standard GUIDE regression tree with *X* as predictors and *Y*_res_ as the response, obtaining predicted residuals *ε*^(*X*). Final predicted probabilities are reconstructed as *p*^ = *p*^*_Z_* + *ε*^(*X*) and the final prediction is truncated to 0 or 1 with a threshold of 0.5.

#### Naive Strategy II: unadjusted GUIDE classification tree

As an unadjusted reference, a GUIDE classification tree is fitted using (*X, Z*) to predict *Y* , omitting the adjustment of confounders.

#### Naive Strategy III: unadjusted GUIDE logistic regression tree

In this approach, a GUIDE logistic regression tree is fitted to (*X, Z*) without adjusting for confounders.

#### scanCT Strategy I: node-specific logistic adjustment

This strategy employs the same GUIDE logistic regression tree, but confounders *Z* are labeled as *f* in the input file to adjust the effect of confounders on leaf nodes. For each terminal node *t*, it fits a logistic regression followed by:

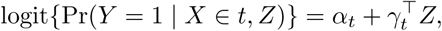

#### scanCT Strategy II: node-specific logistic adjustment assisted by probability

The final strategy is a two-stage hybrid procedure. First, a Random Forest classifier is trained on (*X, Z*) to obtain smoothed probability estimates *p*^_RF_ = Pr(*Y* = 1 *X, Z*). Second, these probabilities are used as a continuous response in a GUIDE logistic regression tree configured as in Strategy I, with *X* for splitting and *Z* as covariates.

#### Evaluation

The tree models were trained on the entire dataset; therefore, the results reported herein represent training performance. To comprehensively evaluate the performance of different strategies, we employed four standard metrics: Precision, Recall, F1-score, and the Area Under the Receiver Operating Characteristic Curve (AUC). Precision measures the proportion of predicted positive cases that are genuinely positive, quantifying the model’s reliability. Recall assesses the model’s ability to identify all actual positive instances from the dataset. The F1-score is the harmonic mean of Precision and Recall, providing a single metric that balances both concerns and is particularly useful for imbalanced datasets. Finally, the AUC represents the degree of separability between classes and is derived from the ROC curve, which plots the True Positive Rate against the False Positive Rate at various threshold settings. These metrics are mathematically defined as follows:

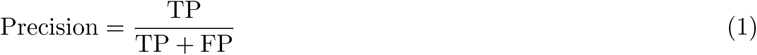

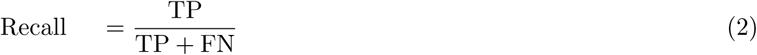

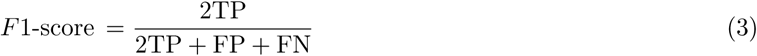

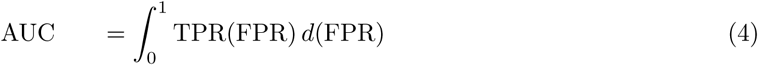

where TP, TN, FP, and FN denote true positives, true negatives, false positives, and false negatives, respectively. TPR refers to the True Positive Rate (Recall), and FPR refers to the False Positive Rate (FP*/*(FP + TN)).

### Correlated features of splitting variables in scanCT I

We performed a comprehensive correlation analysis to identify features significantly associated with the splitting variables selected by the scanCT algorithm. The analytical strategy was determined by the data type of the splitting feature to ensure appropriate statistical testing. For numeric splitting features, we employed a multi-method approach utilizing three distinct statistical tests: the Chi-square test of independence, the Pearson correlation coefficient, and Spearman’s rank correlation coefficient. This triangulation allows us to capture linear, monotonic, and distributional relationships. We identified shared correlated features that consistently ranked among the top candidates across all applicable methods. In cases where the Chi-square test was inapplicable (due to constant numeric features yielding invalid statistics), the consensus set was derived from the intersection of the Pearson and Spearman correlations. From this filtered list, we selected at most the top 10 shared correlated features. For non-numeric splitting features where Pearson and Spearman correlations are undefined, we assessed associations exclusively using the Chi-square test. We selected the top 10 features demonstrating the strongest statistical dependence as indicated by the lowest *p*-values.

### Software Implementation

GUIDE trees were fitted using GUIDE version 43.0 ( http://www.stat.wisc.edu/~loh/treeprogs/scanCT/guideman.pdf). For comparison, CART, Random Forests and XGBoost were implemented in Python using the scikit-learn and xgboost packages.

All Python-based models were initialized with a fixed random seed of 42. Random Forests were implemented using the RandomForestClassifier from the scikit-learn library, configured with 100 trees (n estimators=100). For CART, we utilized the DecisionTreeClassifier from the same package with a maximum depth of 20 and a minimum leaf sample size set to 1% of the total sample size, consistent with the settings used for the GUIDE classification trees. XGBoost was implemented using the XGBClassifier from the xgboost package with the softmax objective (objective=multi:softmax). Unless otherwise specified, all other hyperparameters were kept at their default values.

We applied GUIDE in two configurations:

- **Classification Trees:** We fitted single classification trees (model fitting mode, univariate splits) with interaction tests enabled. We employed estimated class priors, unit misclassification costs, and an exhaustive search for split points. Trees were grown to maximum size without pruning (pruning option 3).
- **Logistic Regression Trees:** We utilized the regression mode with a logistic link function, fitting multiple linear logistic models in each terminal node. Interaction tests were enabled, and trees were unpruned (pruning option 2). To ensure stable coefficient estimation, the minimum terminal node sample size was set to 1% of the sample size.

## Acknowledgements

This work was supported by the National Institutes of Health grant, CA293142 to Y.S. and HG012797 to Y.Z.. Z.L is partially supported by the MD Adnerson Cancer Center PCCSM QS Postbaccalaureate Programs funding.

## Supplementary Materials

### Feature Importance of Random Forests and XGBoost

To ensure a fair comparison with scanCT’s chi-square-based importance, we utilized the standard impurity-based importance metrics provided by Python libraries, which are the default usage in most bioinformatics applications.

- **Random Forest:** Feature importance was calculated using the scikit-learn implementation’s default metric, known as the Mean Decrease in Impurity (MDI) or Gini Importance. This metric measures the total decrease in node impurity (Gini index) weighted by the probability of reaching that node, averaged over all trees in the ensemble.
- **XGBoost:** Feature importance was derived using the feature importances attribute from the

XGBClassifier in the xgboost Python package. This corresponds to the Gain metric, which represents the average gain in accuracy (loss reduction) brought by a feature to the branches it is on.

---

**Table S1.**
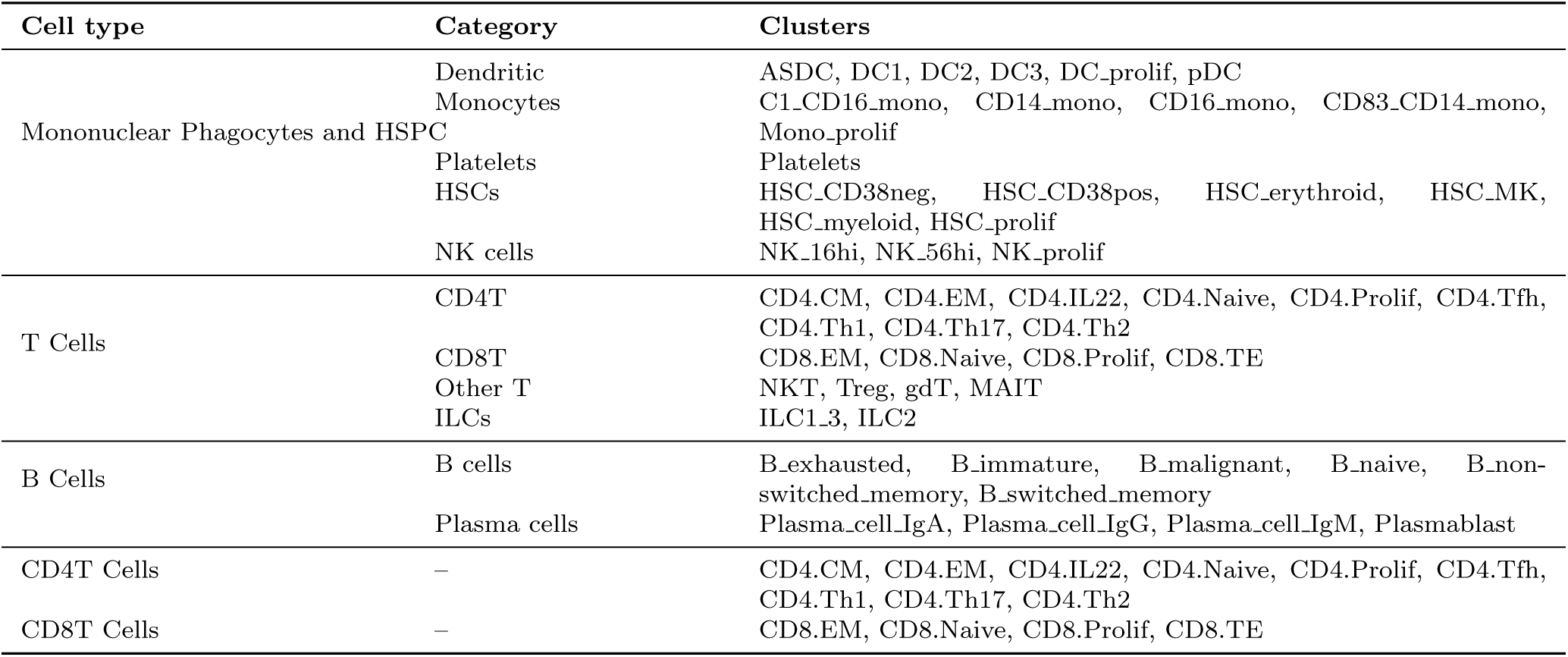
Major immune compartments and included clusters.

**Table S2:**
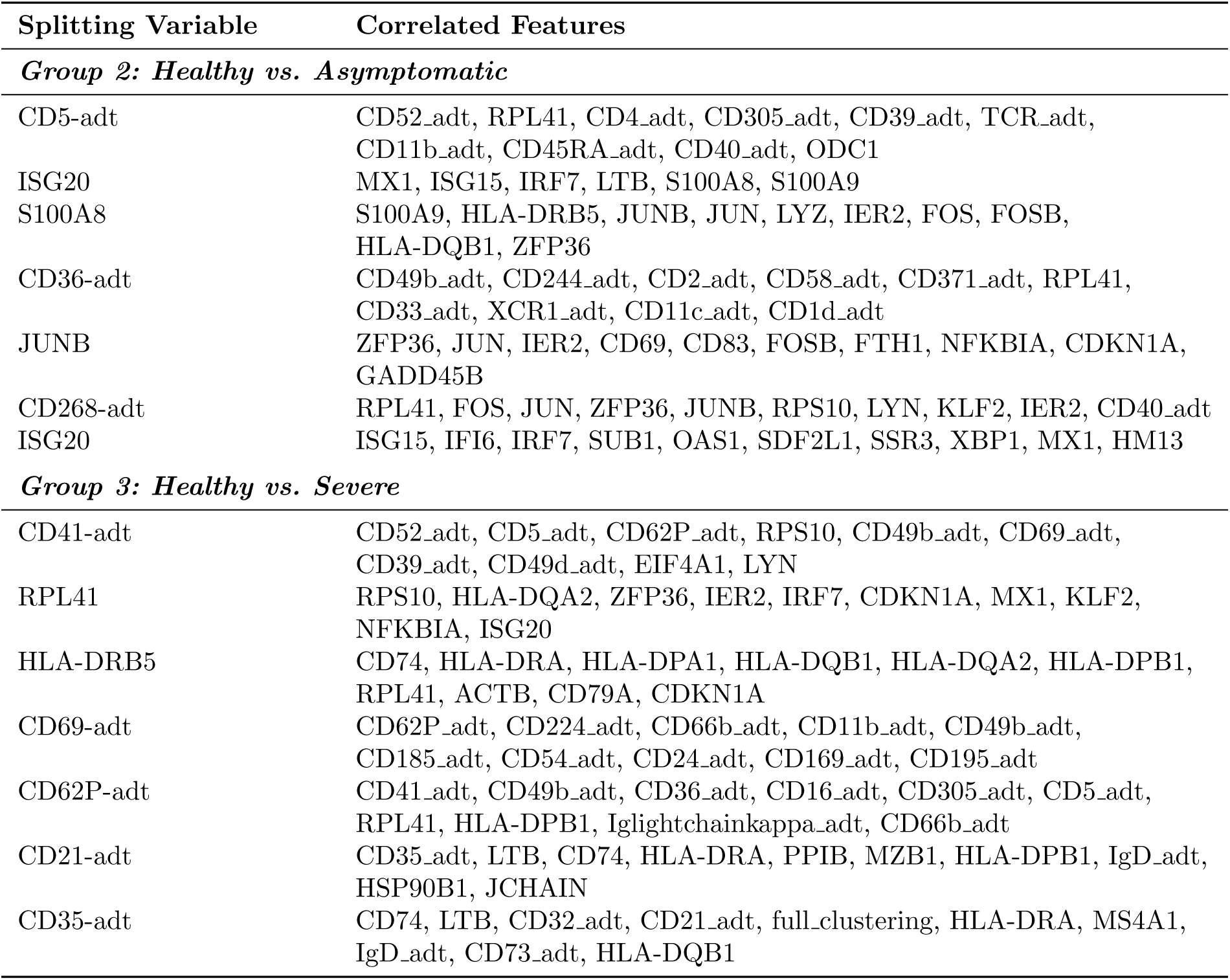
Correlated features in **B Cells**

**Table S3:** Correlated features in **CD4T Cells**

| **Splitting Variable** | **Correlated Features** |
| --- | --- |
| ***Group 2: Healthy vs. Asymptomatic*** |  |
| RPL41 | HBB, CRIP1, SAT1, CALR, ISG15, IFI6, MX1, AC016831.6, KLF2, EIF4A1 |
| CD127-adt | CD27_adt, TCR_adt, CD28_adt, CD26_adt, CD52_adt, CD69_adt, CD95_adt, CD224_adt, HBB, IFITM3 |
| NFKBIA | FOS, JUN, ZFP36, JUNB, IER2, CD69, FOSB, FTH1, DUSP2, KLF2 |
| JUNB | JUN, ZFP36, FOSB, CD69, NFKBIA, DUSP2, KLF2, IER2, FTH1, PTGER4 |
| CD224-adt | CD52_adt, CD45RO_adt, CD41_adt, CD62P_adt, CD66b_adt, CD54_adt, RPL41, CD11b_adt, CRIP1, Ighlightchainkappa_adt |
| Cluster S2 | HSP90B1, SEC11C, FKBP11, CCND2, CD74, TUBA1B, AQP3, MYDGF, HLA-DPA1, PPIB |
| ***Group 3: Healthy vs. Severe*** |  |
| CD2-adt | FOS, COTL1, JUN, JUNB, RPS10, KLF2, IER2, CD69, EIF4A1, CD52_adt |
| CD62P-adt | CD41_adt, CD49b_adt, CD32_adt, CD224_adt, CD66b_adt, Ighlightchainkappa_adt, CD69_adt, CD49f_adt, CD45RO_adt, CD305_adt |
| S100A8 | S100A9, LYZ, JUNB, NFKBIZ, TSHZ2, JUN, CISH, FOS, IER2 |
| SOCS3 | CD69, LDHA, PIM2, NFKBIZ, JUNB, TOB1, FOS, FTH1, NFKBIA, MIF |
| IL7R | S100A4, ACTB, NFKBIA, VIM, FOS, TIMP1, LTB, CD69, S100A11 |
| LDHA | CRIP1, S100A4, ACTB, VIM, IL32, GAPDH, COTL1 |
| MIF | IL32, S100A4, S100A6, GAPDH, CRIP1, VIM, FTH1, S100A11, LGALS1, SEC61B |

**Table S4:** Correlated features in **CD8T Cells**

| **Splitting Variable** | **Correlated Features** |
| --- | --- |
| ***Group 2: Healthy vs. Asymptomatic*** |  |
| RPL41 | HBB, CRIP1, GZMB, FOS, ISG15, IFI6, PRF1, NKG7, RGCC, GZMH |
| CD32-adt | CD35_adt, CD54_adt, CD11b_adt, CD305_adt, CD371_adt, CD226_adt, CD62P_adt, CD11c_adt, Ighlightchainlambda_adt, Ighlightchainkappa_adt |
| CD41-adt | RPL41, KLF2, CD52_adt, CD45RA_adt, CD5_adt, CD27_adt, CD62P_adt, CD305_adt, CD11b_adt, CD32_adt |
| IFI6 | ISG15, ISG20, GZMB, S100A6, HERPUD1, LGALS1, PRF1, S100A4, GZMH, CALR |
| CD32-adt | CD11b_adt, Ighlightchainlambda_adt, CD66b_adt, CD45RO_adt, CD35_adt, CD305_adt, CD52_adt, Ighlightchainkappa_adt, CD169_adt, CD158_adt |
| ***Group 3: Healthy vs. Severe*** |  |
| Clustering in S1 | CCL4, HLA-DPA1, GZMK, HLA-DPB1, SPON2, DUSP2, CCL5, LGALS1, ADGRG1, C12orf75 |
| CD2-adt | RPL41, CD28_adt, KLF2, CD36_adt, CD244_adt, CD27_adt, KLRK1, CD52_adt, CD127_adt, CD122_adt |
| S100A8 | S100A9, LYZ, JUNB, S100A12, JUN, VCAN, NCF2, CLEC7A, CYBB, SPI1 |
| KLRK1 | CCL5, NKG7, GZMK, CD74, LTB, GZMA |
| RPL41 | CRIP1, GZMB, FOS, PRF1, NKG7, RGCC, JUN, KLRK1, CTSD, ZFP36 |
| CD2-adt | KLRK1, RPS10, CD52_adt, CD45RO_adt, CD24_adt, CD244_adt, TCR_adt, CD41_adt, CD7_adt, CD127_adt |
| CD32-adt | CD66b_adt, CD45RO_adt, Ighlightchainlambda_adt, CD35_adt, Ighlightchainkappa_adt, CD54_adt, CD224_adt, CD24_adt, CD11b_adt, CD41_adt |
| GZMB | GNLY, PRF1, NKG7, GZMH, FGFBP2, GZMA, LTB, CST7, S100A4, LGALS1 |

**Table S5:** Correlated features in **Mononuclear cells**

| **Splitting Variable** | **Correlated Features** |
| --- | --- |
| ***Group 2: Healthy vs. Asymptomatic*** |  |
| CD32-adt | HSP90B1, IL1B, S100A8, S100A9, S100A12, IER3, LYZ, HLA-DQA1, HLA-DRA, FKBP11 |
| CD64-adt | IL1B, CCL3, S100A8, S100A9, CXCL8, S100A12, IER3, LYZ, FTL, PLAUR |
| IFITM3 | IFI27, IFI30, RPL41, ISG15, AC245014.3, FCER1G, IER3, MT2A, APOBEC3A, PLAC8 |
| CD52-adt | RPL41, CD305_adt, IFI30, FTL, CD224_adt, Ighlightchainkappa_adt, AC020916.1, CTSS, CD41_adt, AIF1 |
| LST1 | IFI30, MS4A7, SERPINA1, FTH1, LILRB2, CFD, SAT1, CD68, AIF1, PILRA |
| IFI6 | S100A8, S100A9, IFI27, APOBEC3A, S100A12, LYZ, RNASE2, IFITM3, FTL, PLBD1 |
| IFITM3 | APOBEC3A, IFI30, HLA-DRB1, FTL, LST1, SERPINA1, FTH1, FCN1, CFD, SAT1 |
| ***Group 3: Healthy vs. Severe*** |  |
| CD62P-adt | S100A8, S100A9, CCL4, HLA-DRA, IFI30, CST3, HSPA5, AREG, GNLY, FTL |
| Cluster S3 | HSP90B1, CCL3, S100A8, S100A9, CCL4, SEC11C, IFI27, HMOX1, S100A12, LYZ |
| CD32_adt | LYZ, HLA-DRA, CD74, HLA-DRB5, IFI30, CST3, HLA-DPA1, HLA-DRB1, GNLY, FTL |
| MYOM2 | GNLY, AREG, NKG7, IGKV1-12, FCGR3A, GZMA, PRF1, FGFBP2, CX3CR1, ZFP36 |
| GZMB | SPON2, PRF1, NKG7, CST7, GZMA, FGFBP2, GZMH, CTSW |
| KLRK1 | GNLY, NKG7, CTSW, RPS10, GZMA, TRDC, CTSD, CST3, HLA-DQB1, KLRB1 |
| S100A9 | LYZ, FTL, CD14, FCN1, VCAN, MNDA, GRN, TYMP, S100A11, LGALS1 |

**Table S6:** Correlated features in **T Cells**

| **Splitting Variable** | **Correlated Features** |
| --- | --- |
| ***Group 2: Healthy vs. Asymptomatic*** |  |
| RPL41 | HBB, CRIP1, CALR, FOS, ISG15, IFI6, PRF1, NKG7, RGCC, IRF7 |
| CD54-adt | CD11b_adt, CD32_adt, CD305_adt, CD122_adt, CD195_adt, CD66b_adt, CD45RO_adt, CD16_adt, CD69_adt, CD223_adt |
| CD224-adt | RPL41, CD11b_adt, CD52_adt, CD45RO_adt, CD5_adt, CD32_adt, CD69_adt, CD66b_adt, CD35_adt, CD24_adt |
| Cluster S5 | HSP90B1, SEC11C, SDF2L1, FKBP11, CCND2, CD74, TUBA1B, AQP3, MYDGF, SSR3 |
| ***Group 3: Healthy vs. Severe*** |  |
| RPL41 | CRIP1, GZMB, CD8A, FOS, PRF1, NKG7, RGCC, CCL5, GZMH, JUN |
| CD32-adt | CD11b_adt, CD66b_adt, CD35_adt, CD62P_adt, CD41_adt, Ighlightchainlambda_adt, CD52_adt, Ighlightchainkappa_adt, CD2_adt, CD49b_adt |
| CD62P-adt | CD41_adt, CD49b_adt, RPS10, CD52_adt, CD3_adt, CD2_adt, CD49f_adt, CD7_adt, CD24_adt, CD235ab_adt |
| KLRK1 | NKG7, CCL5, CD8A, CTSW, CD8B, CST7, KLRD1, GZMA, GZMH, GNLY |
| CD62P-adt | RPL41, KLF2, CD11b_adt, CD52_adt, CD45RA_adt, CD5_adt, CD32_adt, CD27_adt, CD66b_adt, CD41_adt |
| S100A8 | S100A9, LYZ, JUNB, JUN, RPL41, FOS, NFKBIZ, IER2, ZFP36, CITED2 |
| CD2-adt | RPL41, KLF2, CD52_adt, CD244_adt, CD28_adt, CD45RO_adt, FOS, CD36_adt, CD95_adt, COTL1 |
| SOCS3 | LTB, CCL5, NKG7, PIM2, KLRK1, MAL, CD69, CTSW, SLC40A1, JUNB |
| JUN | FOS, NFKBIA, ZFP36, JUNB, IER2, FOSB, CD69, CCL5, GADD45B, FTH1 |

**Fig. S1.**
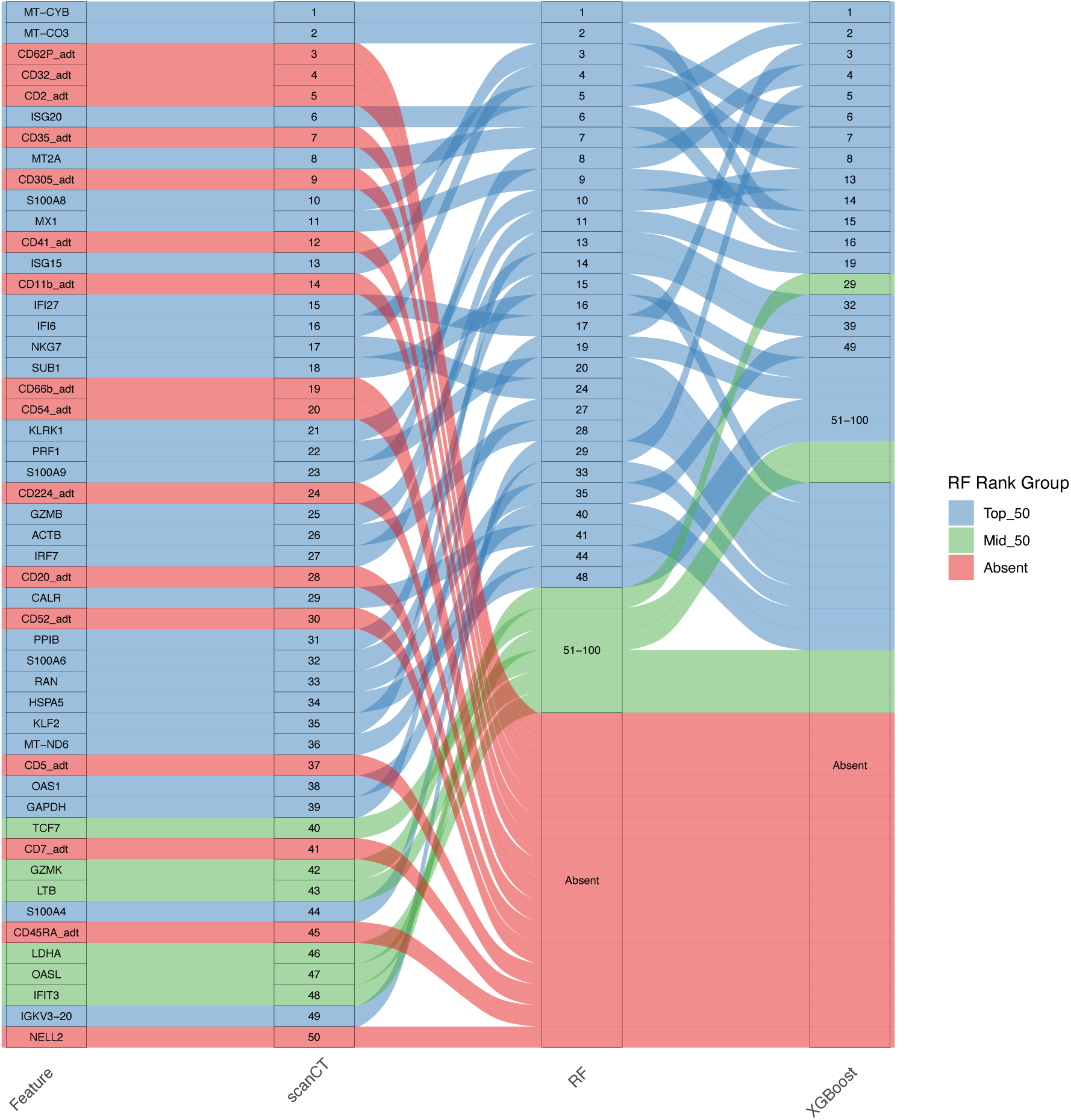
Divergence in Feature Importance Rankings Between scanCT and Ensemble Methods. A Sankey diagram tracing the rank of the top 50 features identified by scanCT (left) to their corresponding positions in Random Forest (RF) and XGBoost (right) on CD8T G1. While a subset of consensus features (blue flows) is present, a significant portion of scanCT’s top features is absent or lowly ranked in RF and XGBoost (red flows). Notably, scanCT prioritizes several antibody-derived tags (ADT), such as *CD64 adt*, *CD56 adt*, and *CD169 adt*. These protein markers are biologically critical for defining immune cell states but are likely penalized in standard impurity-based ensemble methods due to their lower cardinality relative to gene expression features. Green flows indicate features ranked between 51–100 in RF.

**Fig. S2.**
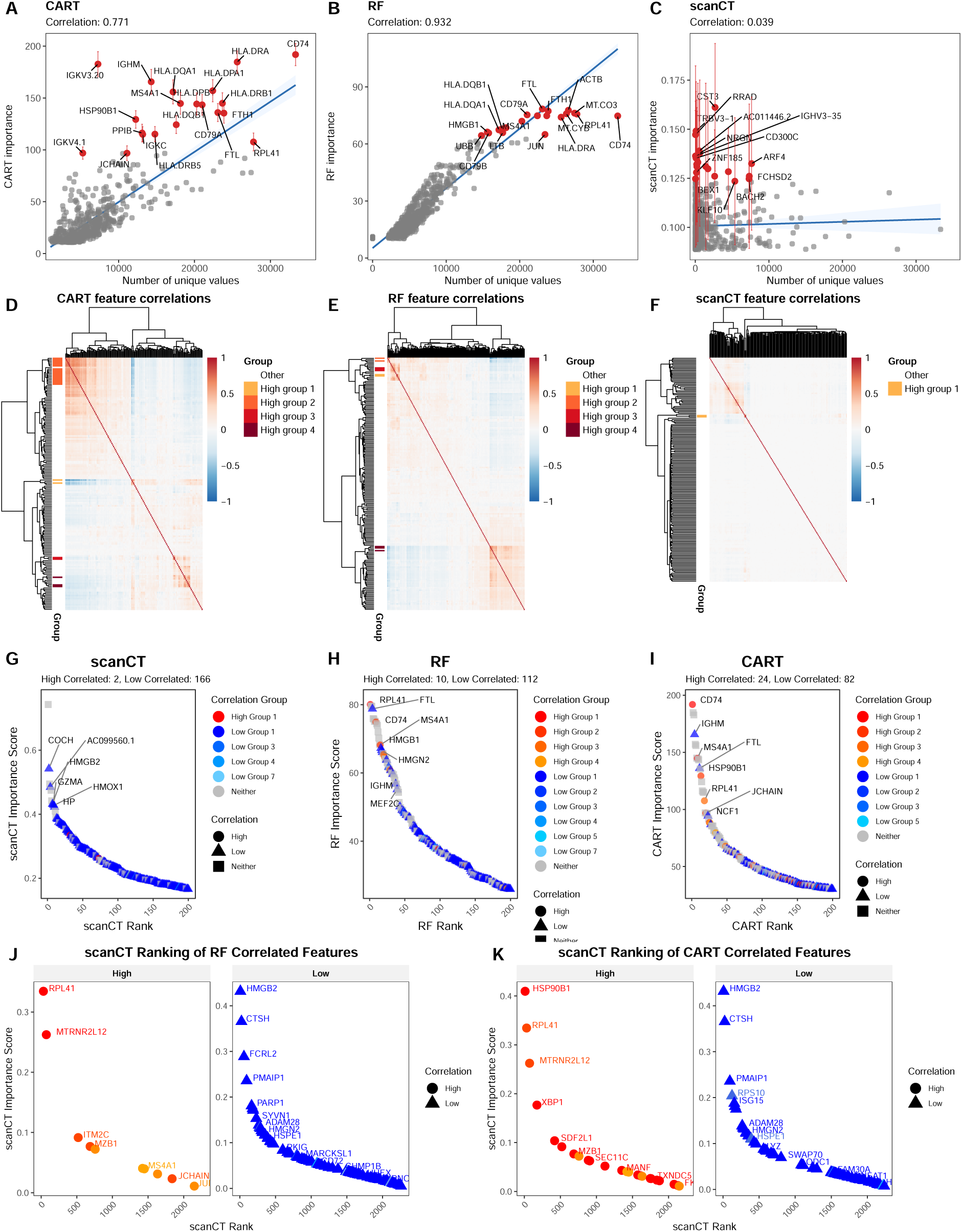
Characterization of feature selection bias in tree-based models. (a–c) Scatter plots of feature importance versus number of unique values for CART (a), Random Forest (RF; b), and scanCT (c) from a permutation experiment on B Group 4 (20 null replicates with *Y* independent of *X*). Blue lines show fitted linear regressions with Pearson correlation coefficients; red points denote the top 20 ranked features with error bars indicating the standard deviation of importance across permutations. **(d–f)** Pairwise Pearson correlation heatmaps for the top 200 features selected by CART (d), RF (e), and scanCT (f), ordered by hierarchical clustering. CART and RF show dense, block-diagonal clusters of strongly correlated predictors, whereas scanCT yields a more diffuse pattern with fewer large, highly correlated blocks. **(g–i)** Rank–abundance plots for the top 200 features selected by CART (g), RF (h), and scanCT (i), stratified by correlation status (highly correlated, low correlated, neither). Highly correlated predictors dominate the upper ranks for CART and RF, whereas low- and uncorrelated features primarily occupy scanCT’s top p_2_o_3_sitions. **(j–k)** Cross-method comparison of scanCT rankings for features belonging to high- and low-correlation groups defined by CART (j) and RF (k). High-correlation groups from CART and RF are dispersed across the scanCT ranking rather than clustering at the top, indicating that scanCT does not inherit the collinearity bias of the comparison methods.

**Fig. S3.**
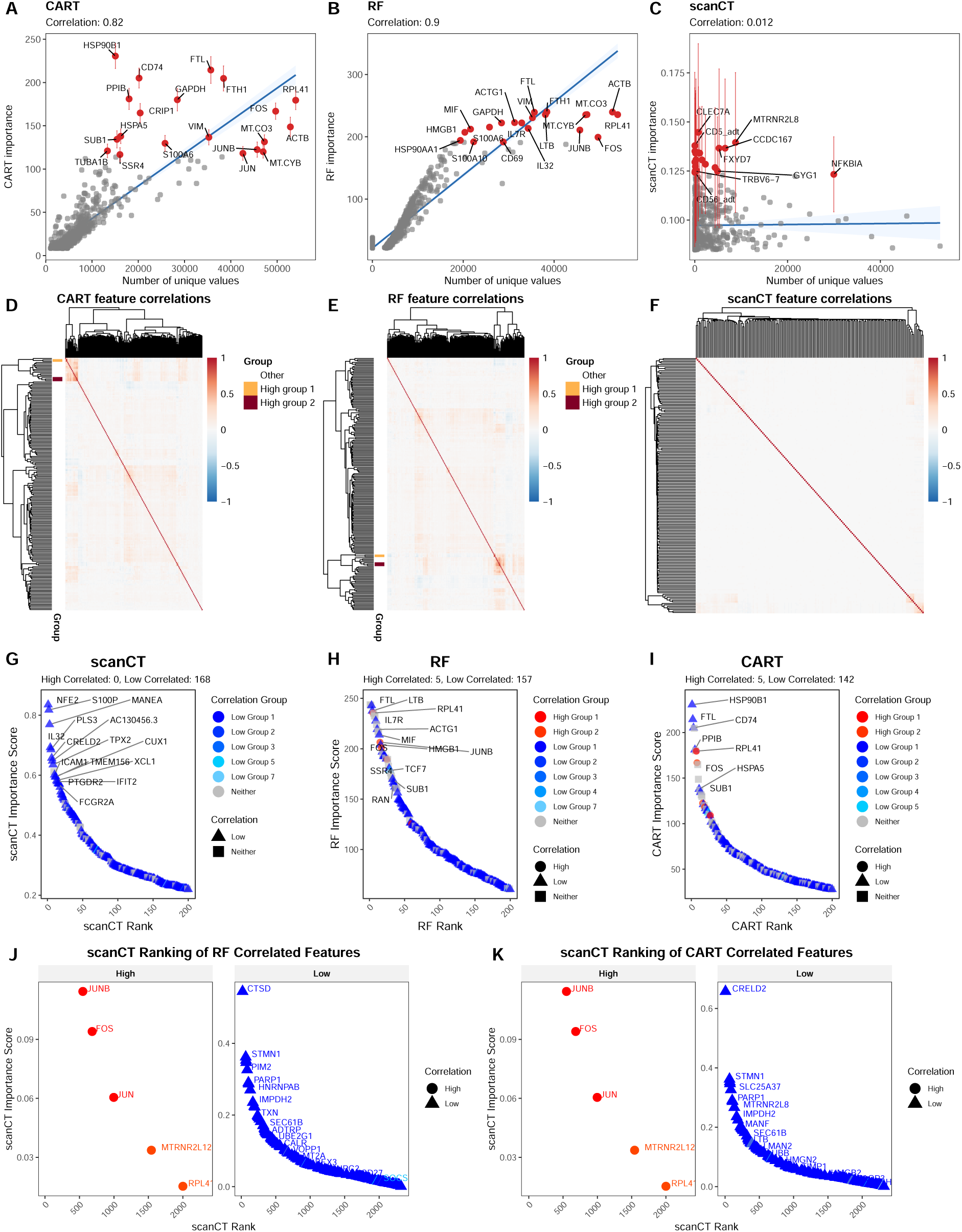
Characterization of feature selection bias in tree-based models. (a–c) Scatter plots of feature importance versus number of unique values for CART (a), Random Forest (RF; b), and scanCT (c) from a permutation experiment on B Group 4 (20 null replicates with Y independent of X). Blue lines show fitted linear regressions with Pearson correlation coefficients; red points denote the top 20 ranked features with error bars indicating the standard deviation of importance across permutations. (d–f) Pairwise Pearson correlation heatmaps for the top 200 features selected by CART (d), RF (e), and scanCT (f), ordered by hierarchical clustering. CART and RF show dense, block-diagonal clusters of strongly correlated predictors, whereas scanCT yields a more diffuse pattern with fewer large, highly correlated blocks. (g–i) Rank–abundance plots for the top 200 features selected by CART (g), RF (h), and scanCT (i), stratified by correlation status (highly correlated, low correlated, neither). Highly correlated predictors dominate the upper ranks for CART and RF, whereas lowand uncorrelated features primarily occupy scanCT’s top positions. (j–k) Cross-method comparison of scanCT rankings for features belonging to high- and low-correlation groups defined by CART (j) and RF (k). High-correlation groups from CART and RF are dispersed across the scanCT ranking rather than clustering at the top, indicating that scanCT does not inherit the collinearity bias of the comparison methods.

**Fig. S4.**
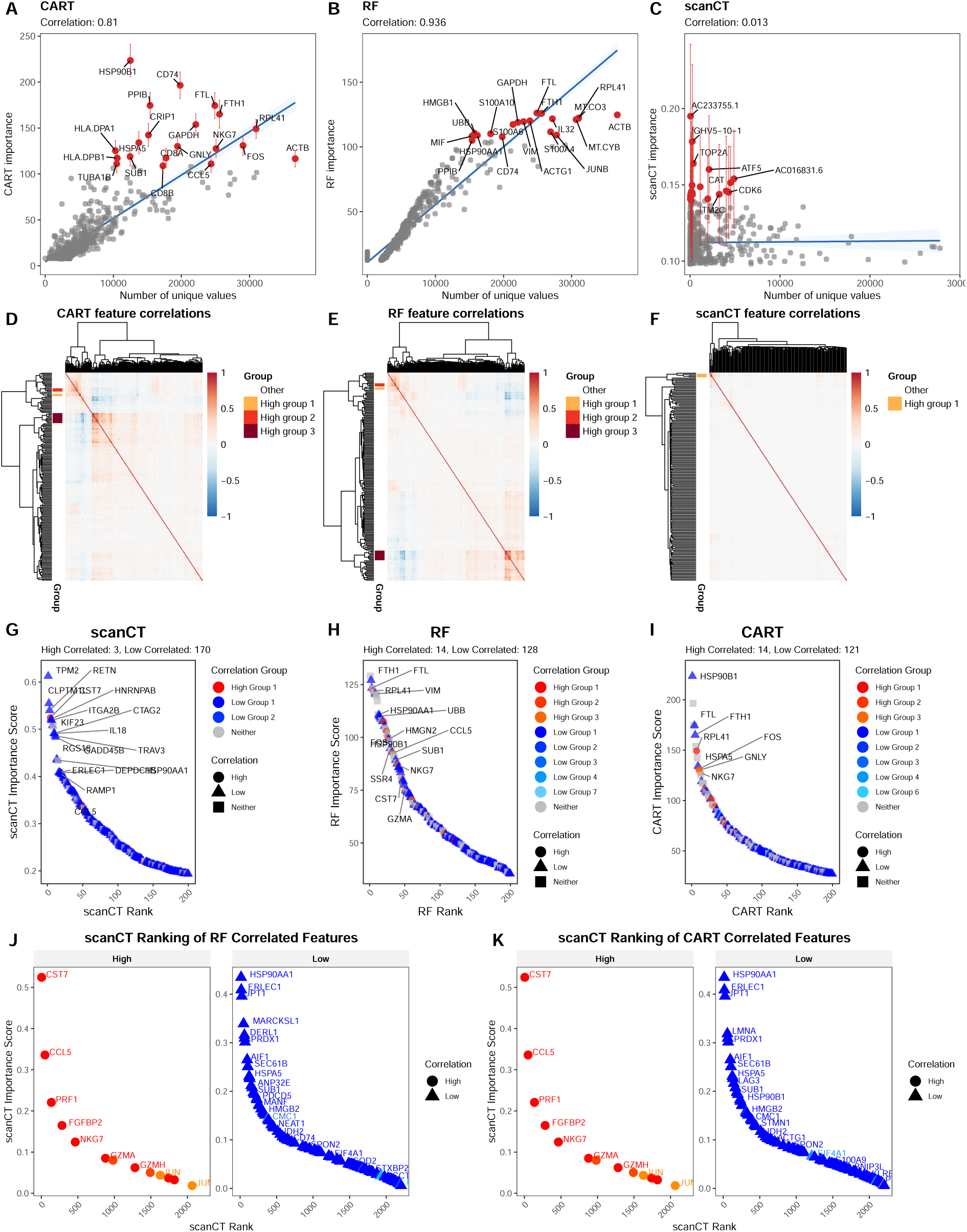
Characterization of feature selection bias in tree-based models. (a–c) Scatter plots of feature importance versus number of unique values for CART (a), Random Forest (RF; b), and scanCT (c) from a permutation experiment on B Group 4 (20 null replicates with Y independent of X). Blue lines show fitted linear regressions with Pearson correlation coefficients; red points denote the top 20 ranked features with error bars indicating the standard deviation of importance across permutations. (d–f) Pairwise Pearson correlation heatmaps for the top 200 features selected by CART (d), RF (e), and scanCT (f), ordered by hierarchical clustering. CART and RF show dense, block-diagonal clusters of strongly correlated predictors, whereas scanCT yields a more diffuse pattern with fewer large, highly correlated blocks. (g–i) Rank–abundance plots for the top 200 features selected by CART (g), RF (h), and scanCT (i), stratified by correlation status (highly correlated, low correlated, neither). Highly correlated predictors dominate the upper ranks for CART and RF, whereas lowand uncorrelated features primarily occupy scanCT’s top positions. (j–k) Cross-method comparison of scanCT rankings for features belonging to high- and low-correlation groups defined by CART (j) and RF (k). High-correlation groups from CART and RF are dispersed across the scanCT ranking rather than clustering at the top, indicating that scanCT does not inherit the collinearity bias of the comparison methods.

**Fig. S5.**
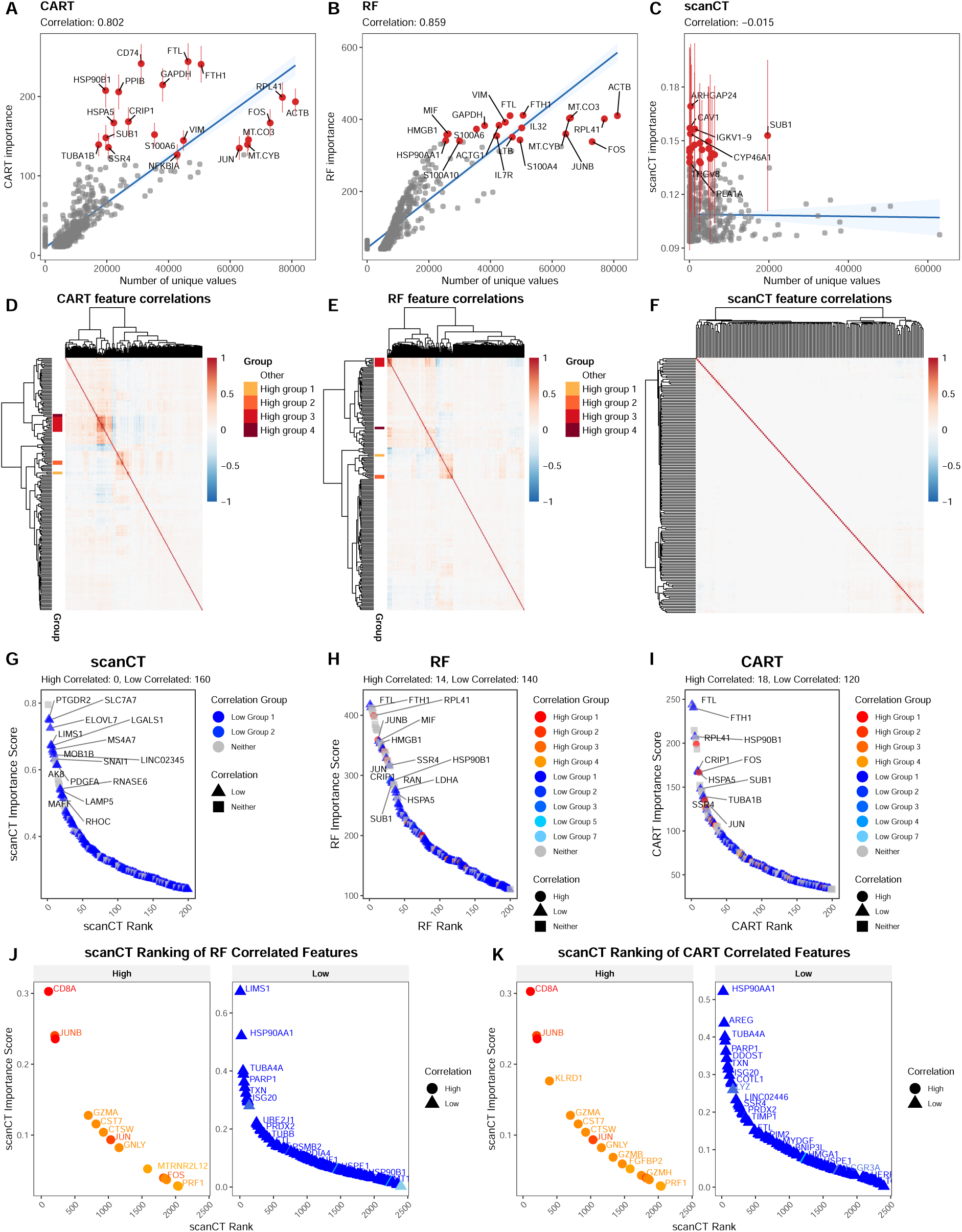
Characterization of feature selection bias in tree-based models. (a–c) Scatter plots of feature importance versus number of unique values for CART (a), Random Forest (RF; b), and scanCT (c) from a permutation experiment on B Group 4 (20 null replicates with Y independent of X). Blue lines show fitted linear regressions with Pearson correlation coefficients; red points denote the top 20 ranked features with error bars indicating the standard deviation of importance across permutations. (d–f) Pairwise Pearson correlation heatmaps for the top 200 features selected by CART (d), RF (e), and scanCT (f), ordered by hierarchical clustering. CART and RF show dense, block-diagonal clusters of strongly correlated predictors, whereas scanCT yields a more diffuse pattern with fewer large, highly correlated blocks. (g–i) Rank–abundance plots for the top 200 features selected by CART (g), RF (h), and scanCT (i), stratified by correlation status (highly correlated, low correlated, neither). Highly correlated predictors dominate the upper ranks for CART and RF, whereas lowand uncorrelated features primarily occupy scanCT’s top positions. (j–k) Cross-method comparison of scanCT rankings for features belonging to high- and low-correlation groups defined by CART (j) and RF (k). High-correlation groups from CART and RF are dispersed across the scanCT ranking rather than clustering at the top, indicating that scanCT does not inherit the collinearity bias of the comparison methods.

**Fig. S6.**
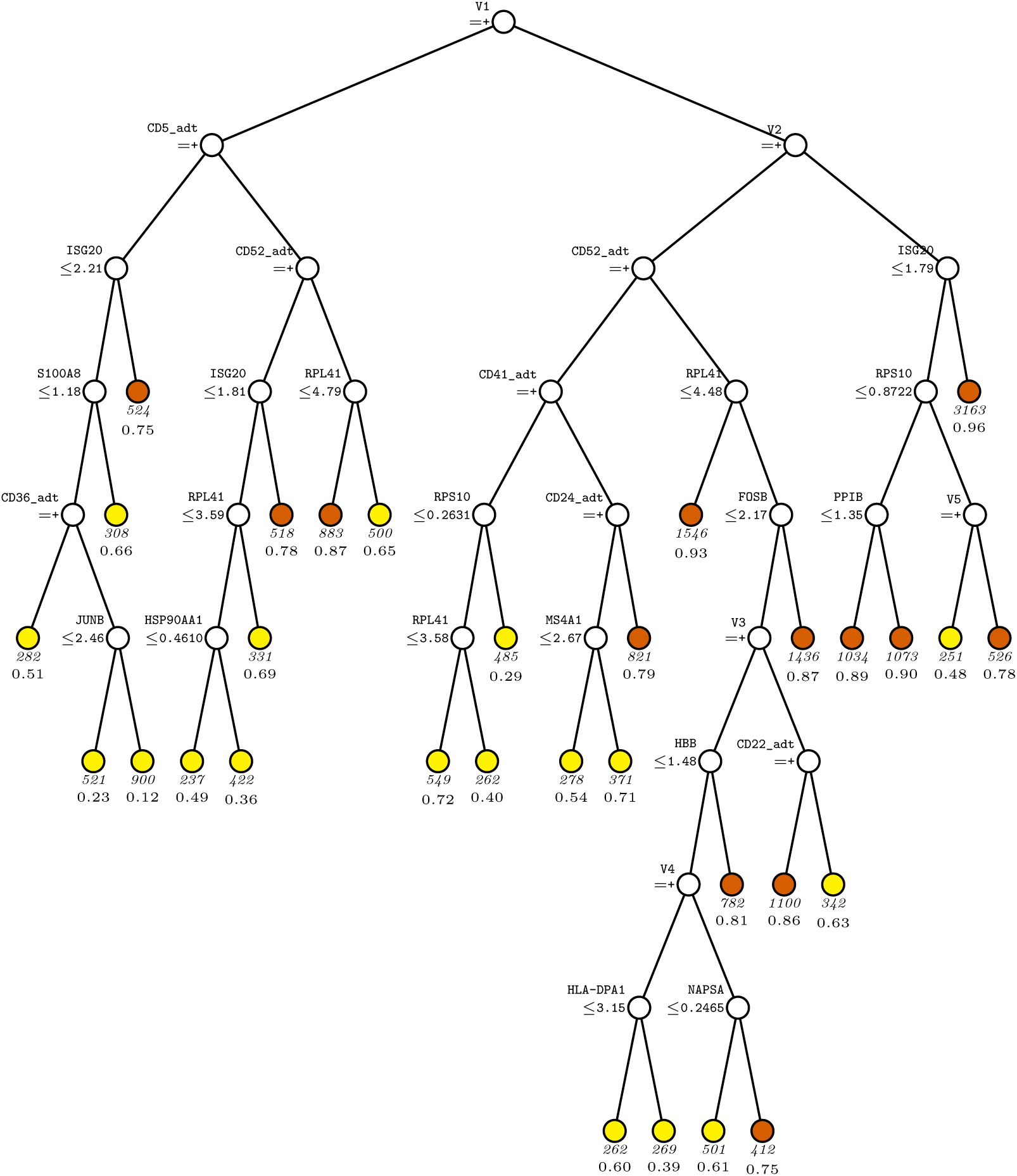
scanCT I tree for predicting. *Y* **in B-cell G2**. At each split, an observation goes to the left branch if and only if the condition is satisfied.Splitting variables are denoted by *V_i_* with the mapping: *V*_1_=CD62P adt; *V*_2_=CD268 adt; *V*3=CD45RA adt; *V*4=CD185 adt; *V*5=CD305 adt. Sample size (in italics) and proportion of *Y* = 1 are shown below each terminal node. Terminal nodes with proportions of 1s above and below value of 0.74 at root node mean are painted vermillion and yellow, respectively.

**Fig. S7.**
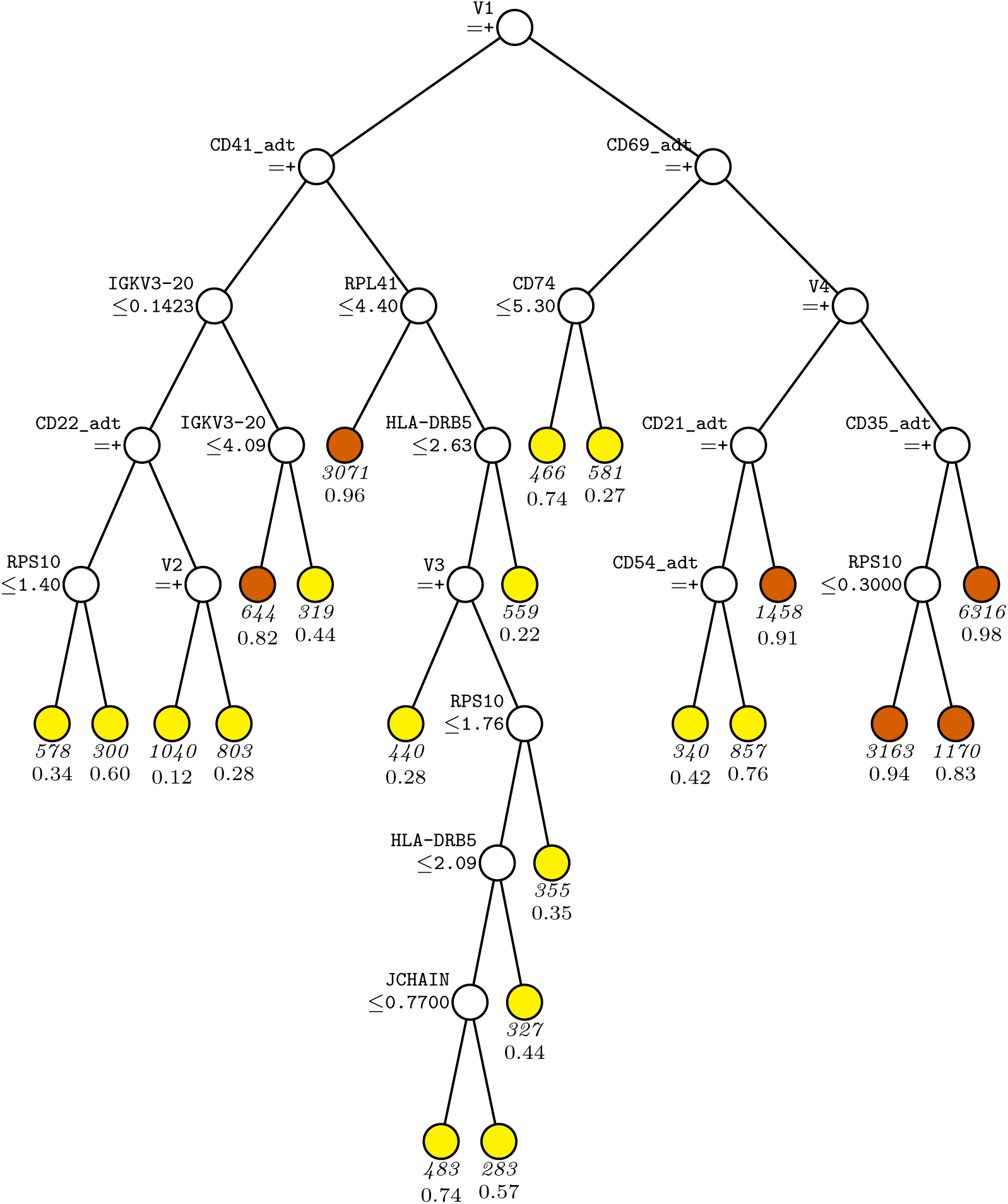
scanCT I tree for predicting. *Y* **in B-cell G3**. At each split, an observation goes to the left branch if and only if the condition is satisfied. Splitting variables are denoted by *V_i_* with the mapping: *V*_1_=CD268 adt; *V*_2_=CD62P adt; *V*_3_=CD305 adt; *V*4=CD62P adt. Sample size (in italics) and proportion of *Y* = 1 are shown below each terminal node. Terminal nodes with proportions of 1s above and below value of 0.77 at root node mean are painted vermillion and yellow, respectively.

**Fig. S8.**
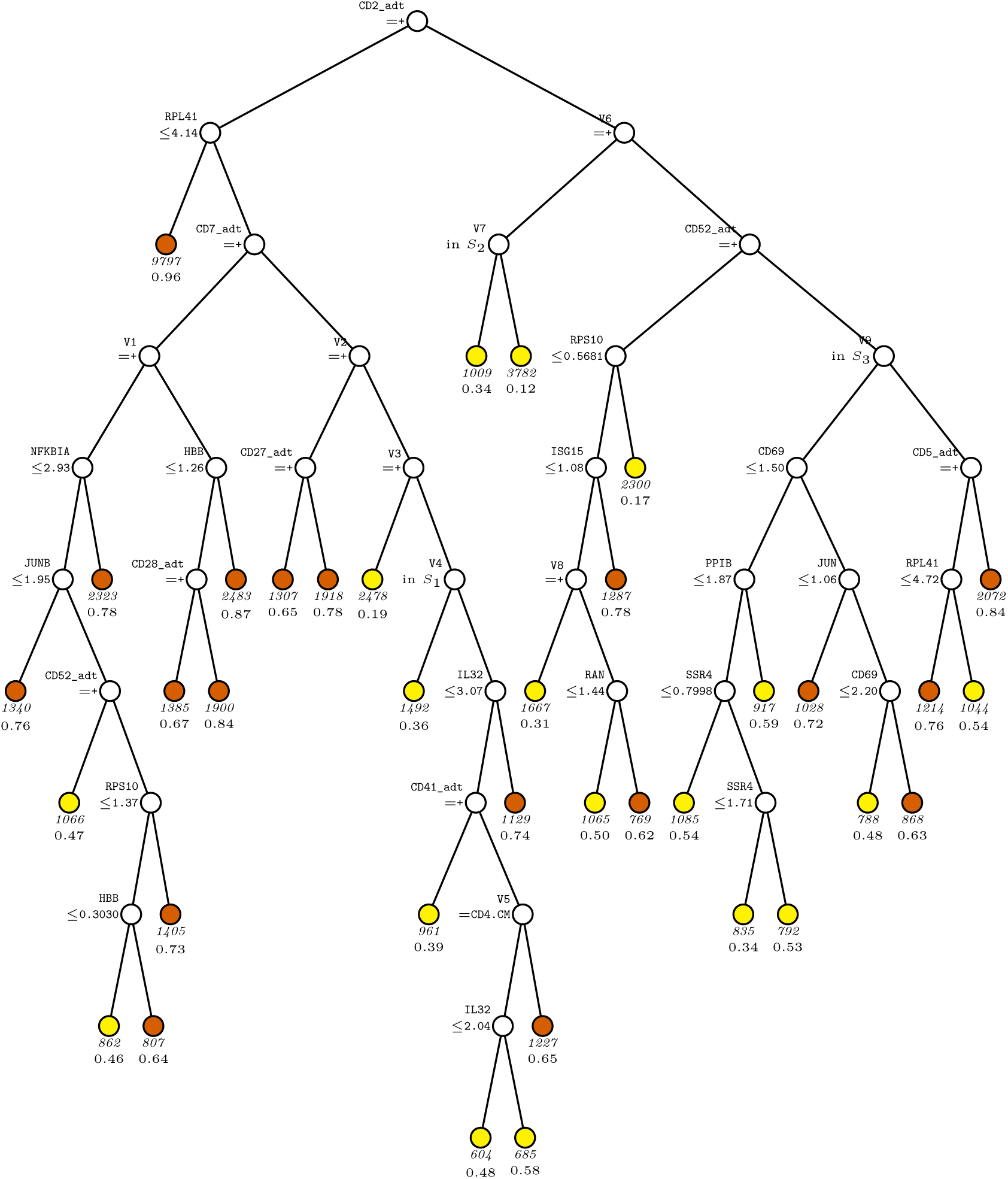
scanCT I tree for predicting. *Y* **in CD4T-cell G2**.At each split, an observation goes to the left branch if and only if the condition is satisfied. Splitting variables are denoted by *V_i_* with the mapping: *V*_1_=CD127 adt; *V*_2_=CD127 adt; *V*3=CD169 adt; *V*4=full clustering; *V*5=full clustering; *V*6=CD224 adt; *V*7=full clustering; *V*8=CD62P adt; *V*9=full clustering. *S*1={CD4.IL22, CD4.Th17}; *S*2={CD4.CM, CD4.EM, CD4.Prolif, CD4.Tfh}; *S*3={CD4.IL22, CD4.Naive, CD4.Th2}. Sample size (in italics) and proportion of *Y* = 1 are shown below each terminal node. Terminal nodes with proportions of 1s above and below value of 0.62 at root node mean are painted vermillion and yellow, respectively.

**Fig. S9.**
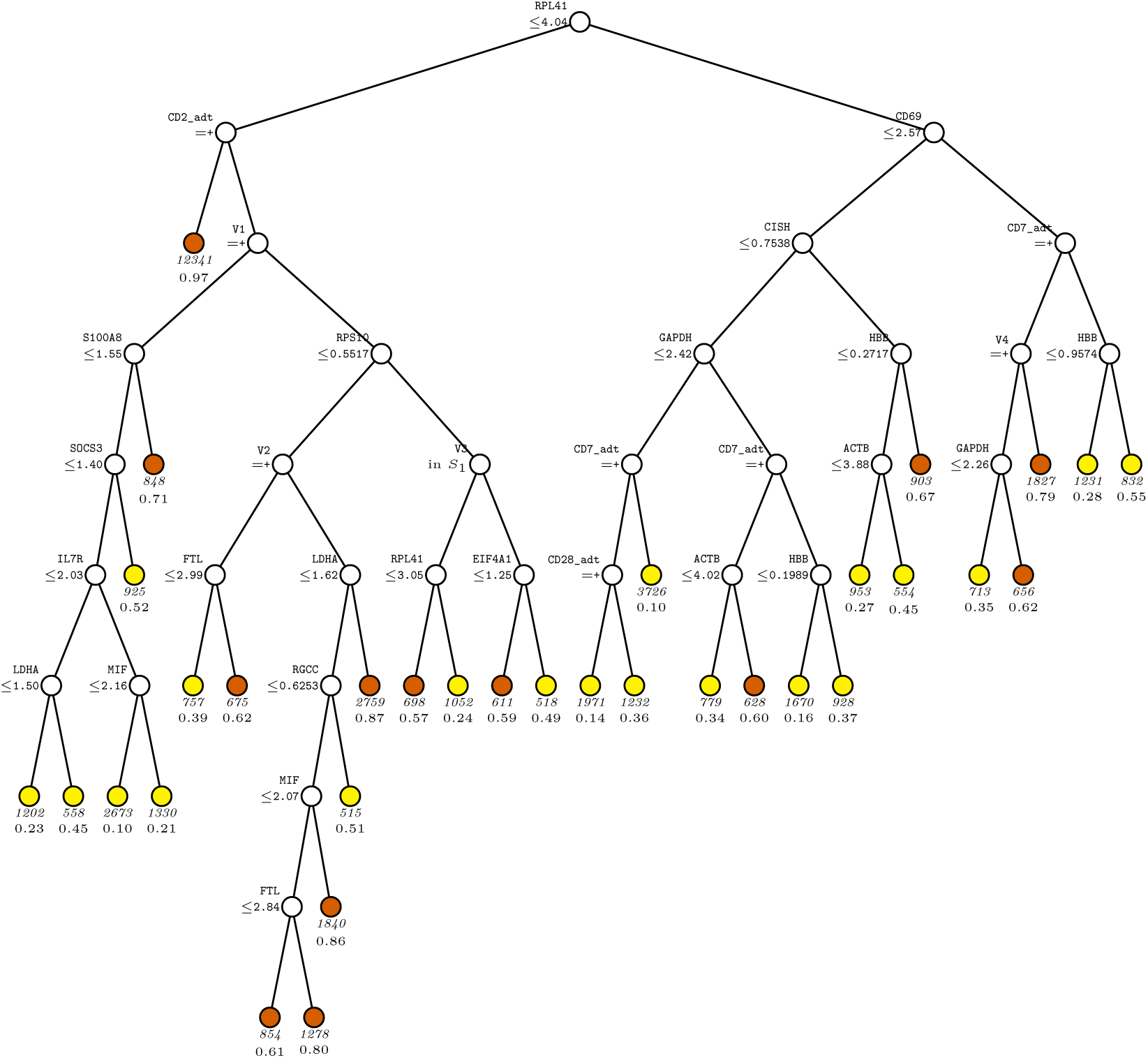
scanCT I tree for predicting. *Y* **in CD4T-cell G3**.At each split, an observation goes to the left branch if and only if the condition is satisfied. Splitting variables are denoted by *V_i_* with the mapping: *V*_1_=CD62P adt; *V*_2_=CD224 adt; *V*3=full clustering; *V*4=CD127 adt. *S*1={CD4.Naive, CD4.Th2}. Sample size (in italics) and proportion of *Y* = 1 are shown below each terminal node. Terminal nodes with proportions of 1s above and below value of 0.56 at root node mean are painted vermillion and yellow, respectively.

**Fig. S10.**
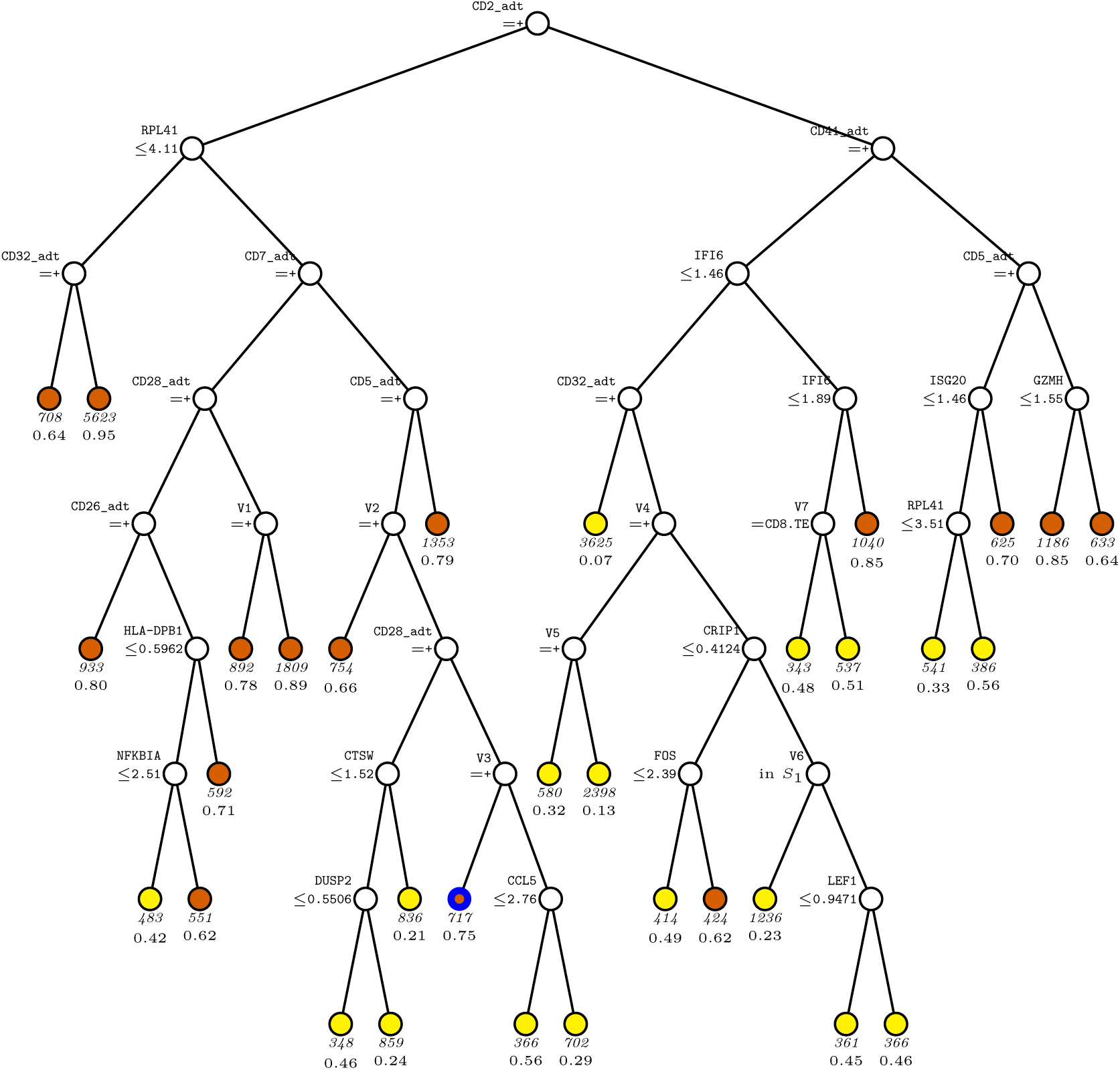
scanCT I tree for predicting. *Y* **in CD8T-cell G2.** At each split, an observation goes to the left branch if and only if the condition is satisfied. Splitting variables are denoted by *V_i_*, defined as: *V*_1_*, V*_2_=CD127 adt; *V*_3_=CD244 adt; *V*4=CD62P adt; *V*5=CD49b adt; *V*6*, V*7=full clustering. Set *S*1={CD8.EM, CD8.TE}. Below each terminal node, the sample size is shown in italics, followed by the proportion of *Y* = 1. Terminal nodes with proportions of 1s above and below value of 0.57 at root node are painted vermillion and yellow respectively. **The node outlined in blue indicates the subgroup selected for node analysis.**

**Fig. S11.**
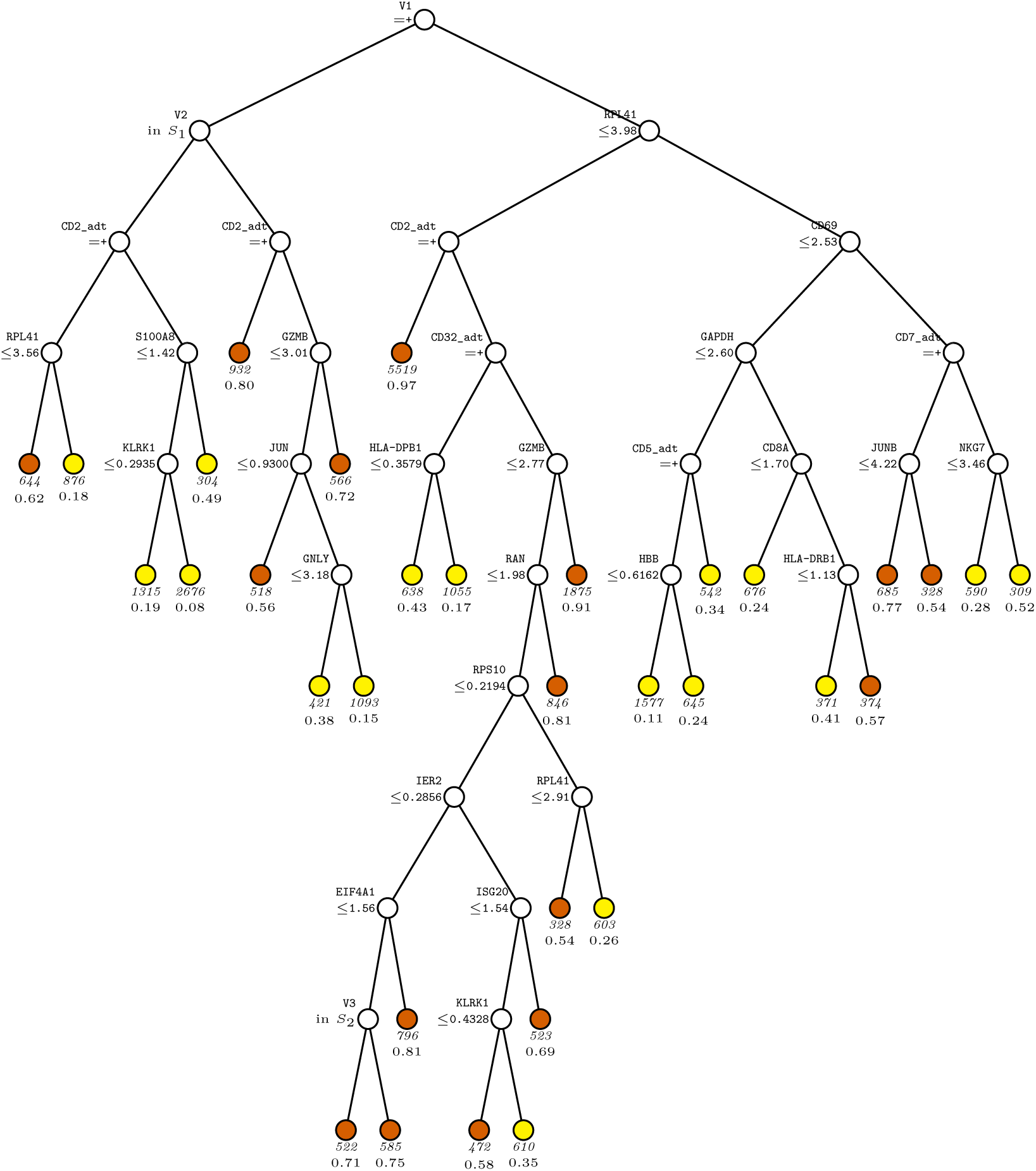
scanCT I tree for predicting. *Y* **in CD8T-cell G3**. At each split, an observation goes to the left branch if and only if the condition is satisfied. Splitting variables are denoted by *V_i_* with the mapping: *V*_1_=CD62P adt; *V*_2_=full clustering; *V*3=full clustering. *S*1={CD8.EM, CD8.Naive}; *S*2={CD8.Naive}. Sample size (in italics) and proportion of *Y* = 1 are shown below each terminal node. Terminal nodes with proportions of 1s above and below value of 0.53 at root node are painted vermillion and yellow respectively.

**Fig. S12.**
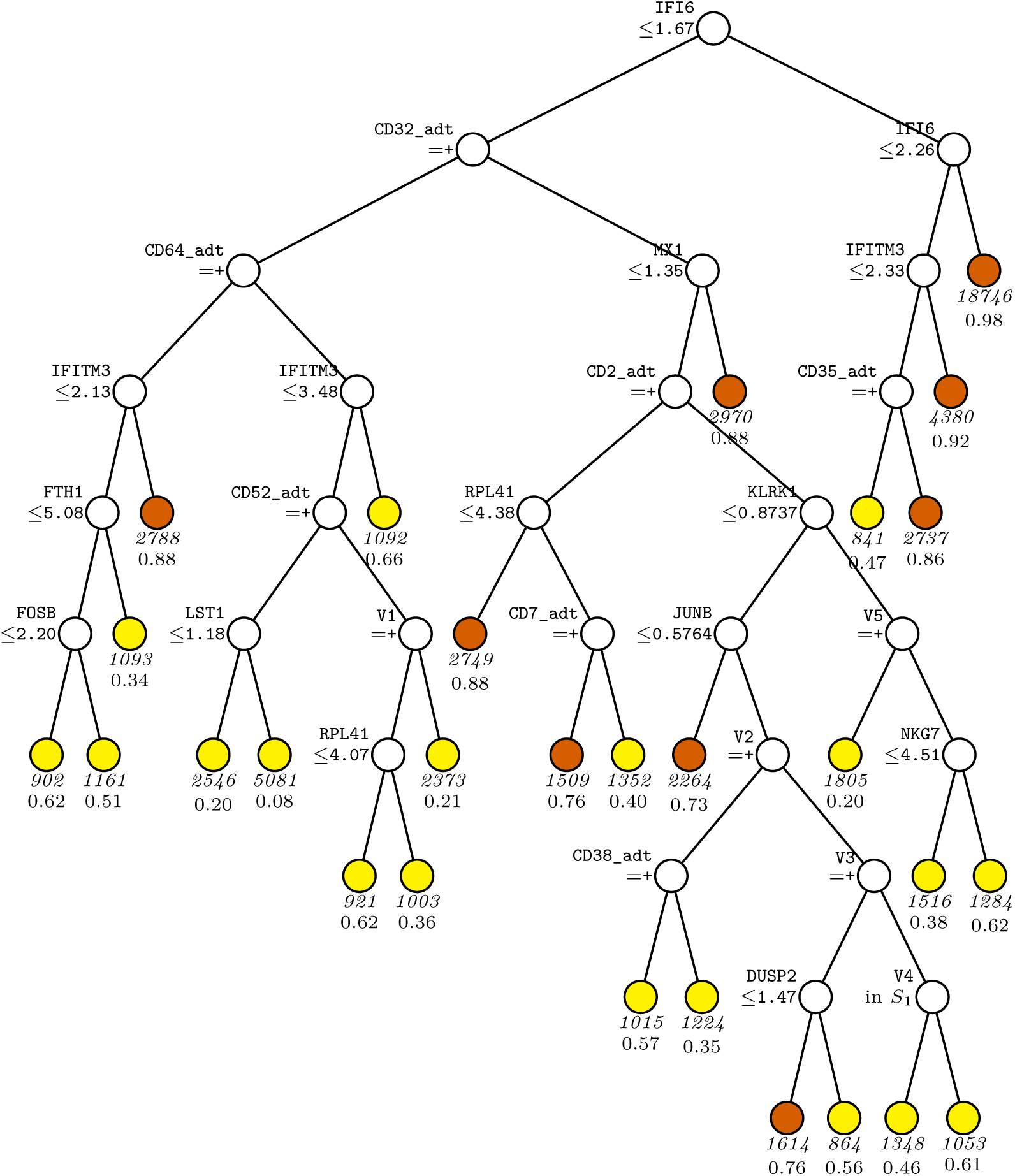
scanCT I tree for predicting. *Y* **in Mononuclear-cell G2**.At each split, an observation goes to the left branch if and only if the condition is satisfied. Splitting variables are denoted by *V_i_* with the map- ping: *V*1=CD244 adt; *V*2=CD11b adt; *V*3=CD244 adt; *V*4=full clustering; *V*5=CD62P adt. *S*1={CD16 mono, DC1, DC2, DC3, DC prolif, HSC erythroid, HSC myeloid, NK 16hi}. Sample size (in italics) and proportion of *Y* = 1 are shown below each terminal node. Terminal nodes with proportions of 1s above and below value of 0.68 at root node are painted vermillion and yellow respectively.

**Fig. S13.**
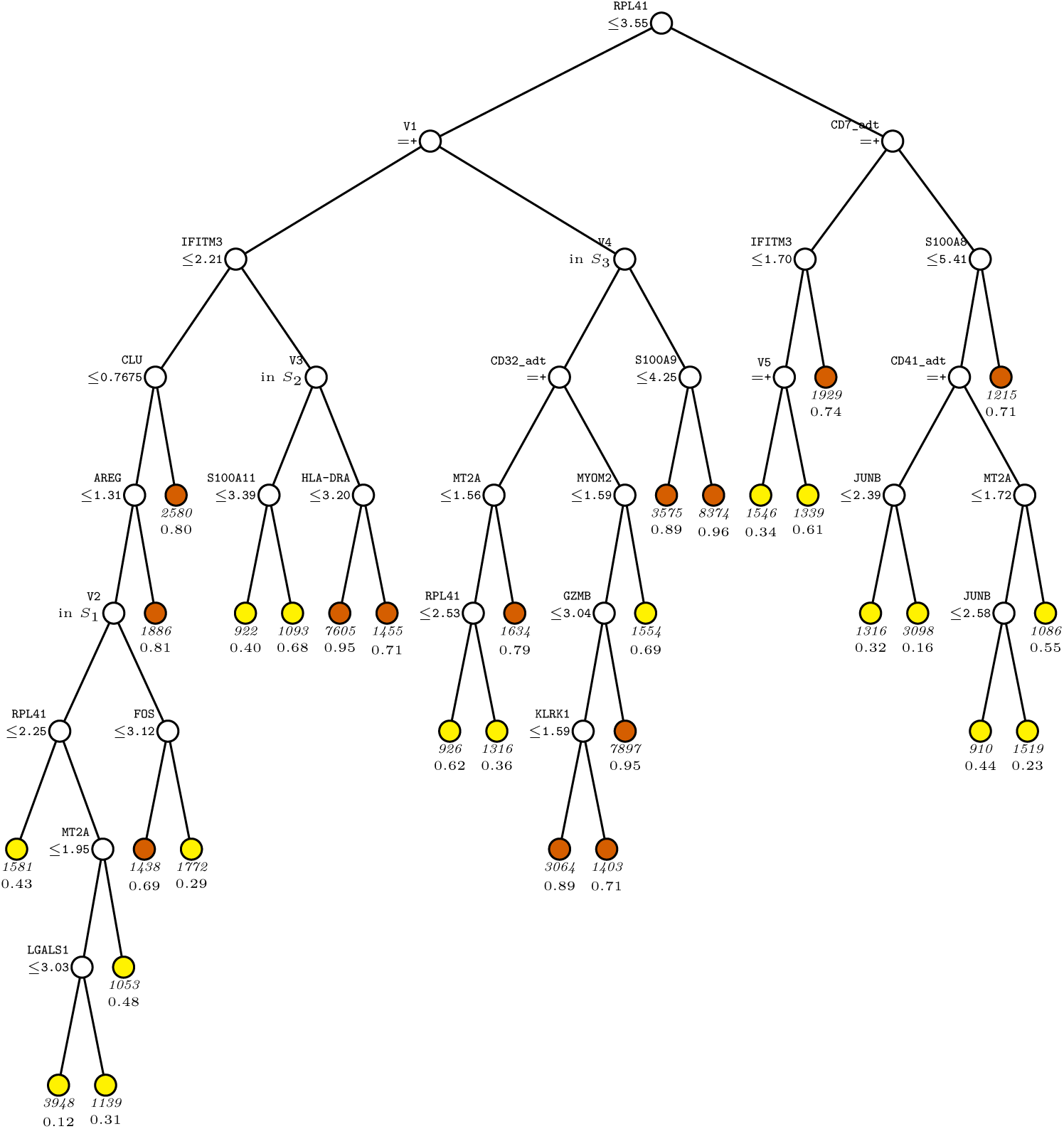
scanCT I tree for predicting. *Y* **in Mononuclear-cell G3**. At each split, an observation goes to the left branch if and only if the condition is satisfied. Splitting variables are denoted by *V_i_* with the mapping: *V*_1_=CD62P adt; *V*2=full clustering; *V*3=full clustering; *V*4=full clustering; *V*5=CD328 adt. *S*1={ASDC, C1 CD16 mono, CD16 mono, CD83 CD14 mono, DC1, DC2, DC3, NK 16hi, NK 56hi}; *S*2={CD16 mono, DC2, NK 56hi}; *S*3={ASDC, CD16 mono, DC1, DC2, DC3, DC prolif, HSC erythroid, NK 16hi, NK 56hi, pDC}. Sample size (in italics) and proportion of *Y* = 1 are shown below each terminal node. Terminal nodes with proportions of 1s above and below value of 0.69 at root node are painted vermillion and yellow respectively.

**Fig. S14.**
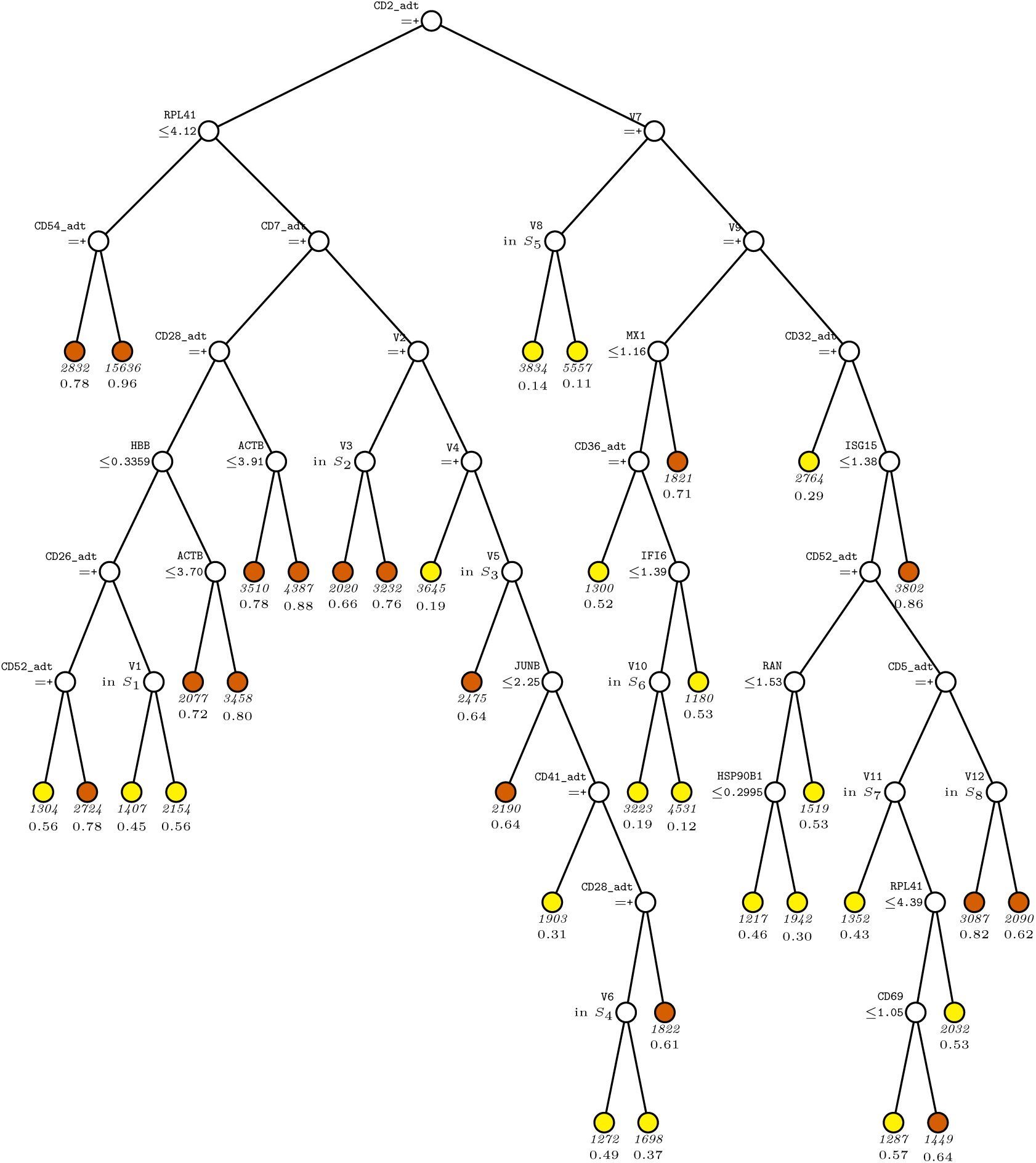
scanCT I tree for predicting. *Y* **in T-cell G2**. At each split, an observation goes to the left branch if and only if the condition is satisfied. Splitting variables are denoted by *V_i_* with the mapping: *V*_1_=full clustering; *V*_2_=CD127 adt; *V*3=full clustering; *V*4=CD169 adt; *V*5=full clustering; *V*6=full clustering; *V*7=CD224 adt; *V*8=full clustering; *V*9=CD62P adt; *V*10=full clustering; *V*11=full clustering; *V*12=full clustering. *S*1={CD4.Naive, CD8.Naive, ILC1 3}; *S*2={CD4.IL22, CD4.Th2, CD8.EM, CD8.TE, MAIT}; *S*3={CD4.EM, CD4.Prolif, CD4.Tfh, CD4.Th2, CD8.Prolif, CD8.TE, NKT, Treg}; *S*4={CD4.CM, CD4.Naive, CD4.Th1, ILC1 3, ILC2}; *S*5={CD4.CM, CD4.IL22, CD4.Th1, CD8.Naive, CD8.TE, ILC1 3, Treg}; *S*6={CD4.IL22, CD4.Naive, ILC2, MAIT, NKT}; *S*7={CD4.IL22, CD4.Th2, CD8.EM, CD8.TE, MAIT, NKT, Treg}; *S*8={CD4.CM, CD4.EM, CD4.IL22, CD4.Prolif, CD4.Tfh, CD4.Th1, CD8.Naive, CD8.Prolif, gdT}. Sample size (in italics) and proportion of *Y* = 1 are shown below each terminal node. Terminal nodes with proportions of 1s above and below value of 0.59 at root node are painted vermillion and yellow respectively.

**Fig. S15.**
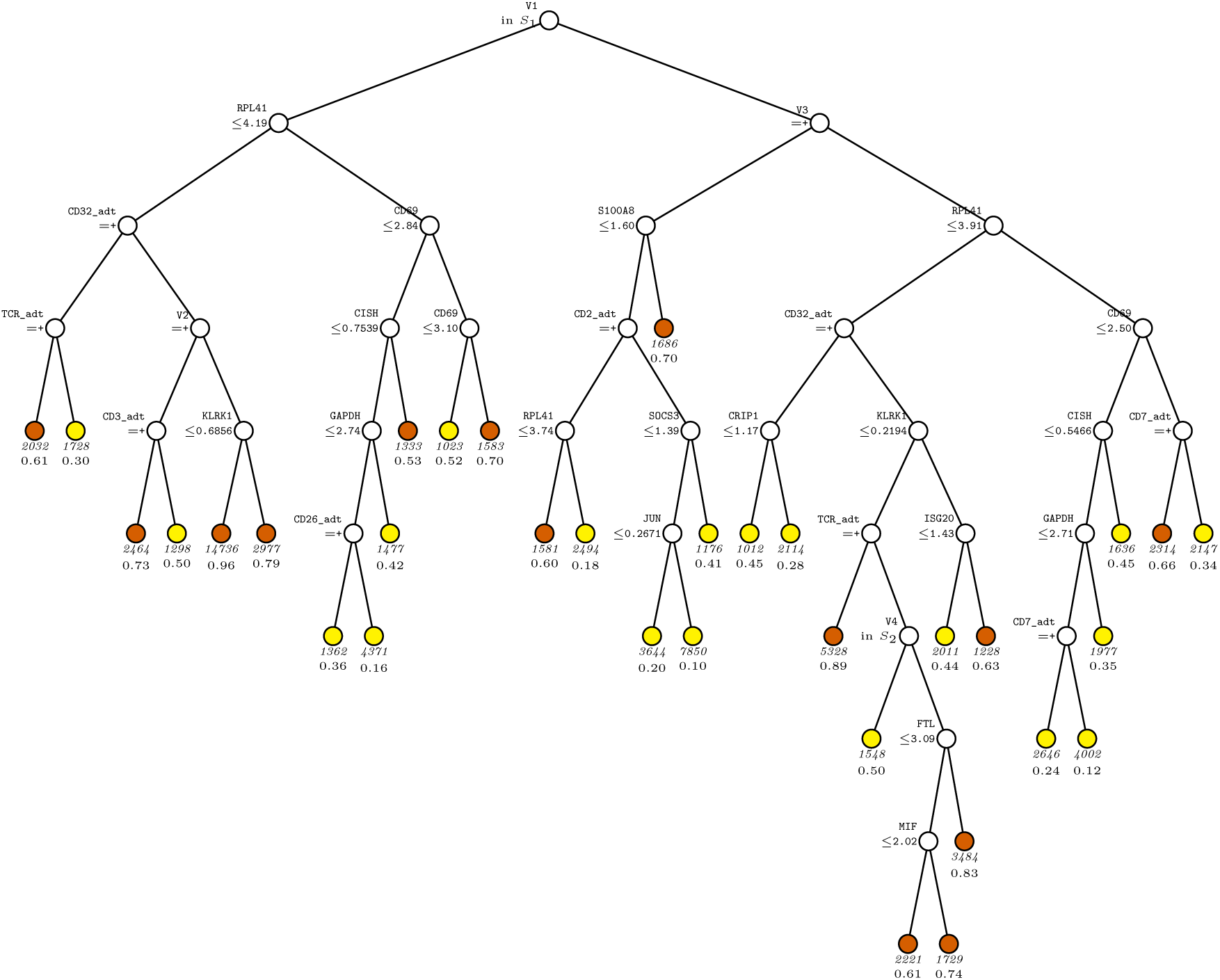
scanCT I tree for predicting. *Y* **in T-cell G3**. At each split, an observation goes to the left branch if and only if the condition is satisfied. Splitting variables are denoted by *V_i_* with the mapping: *V*_1_=full clustering; *V*_2_=CD62P adt; *V*3=CD62P adt; *V*4=full clustering. *S*1={CD4.CM, CD4.EM, CD4.Prolif, CD4.Tfh, CD4.Th17, CD8.Prolif, CD8.TE}; *S*2={CD4.IL22, CD4.Th2, MAIT, Treg}. Sample size (in italics) and proportion of *Y* = 1 are shown below each terminal node. Terminal nodes with proportions of 1s above and below value of 0.53 at root node are painted vermillion and yellow respectively.

**Fig. S16.**
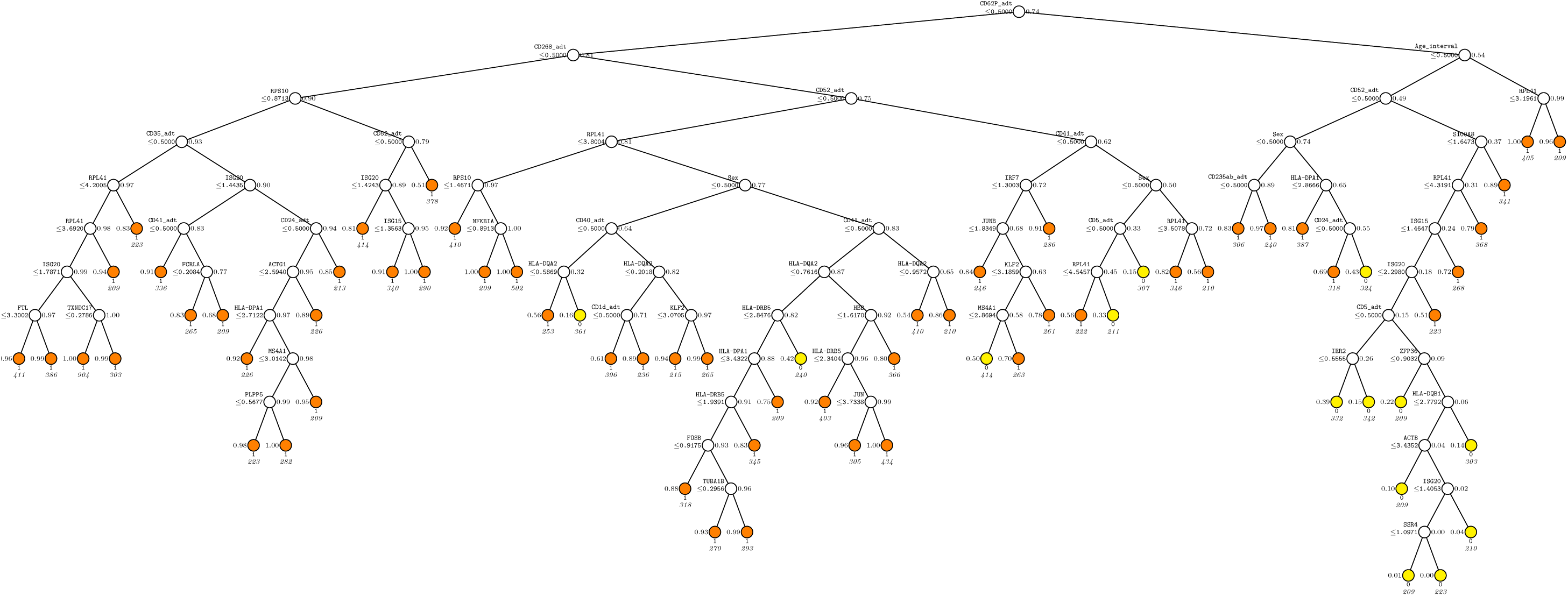
CART tree in B-cell G2. At each split, an observation goes to the left branch if and only if the condition is satisfied. Predicted class and sample size (in italics) printed below each terminal nodes; class sample proportion for Y = 1 beside node. Terminal nodes with proportions of 1s above and below value of 0.5 are painted vermillion and yellow respectively.

**Fig. S17.**
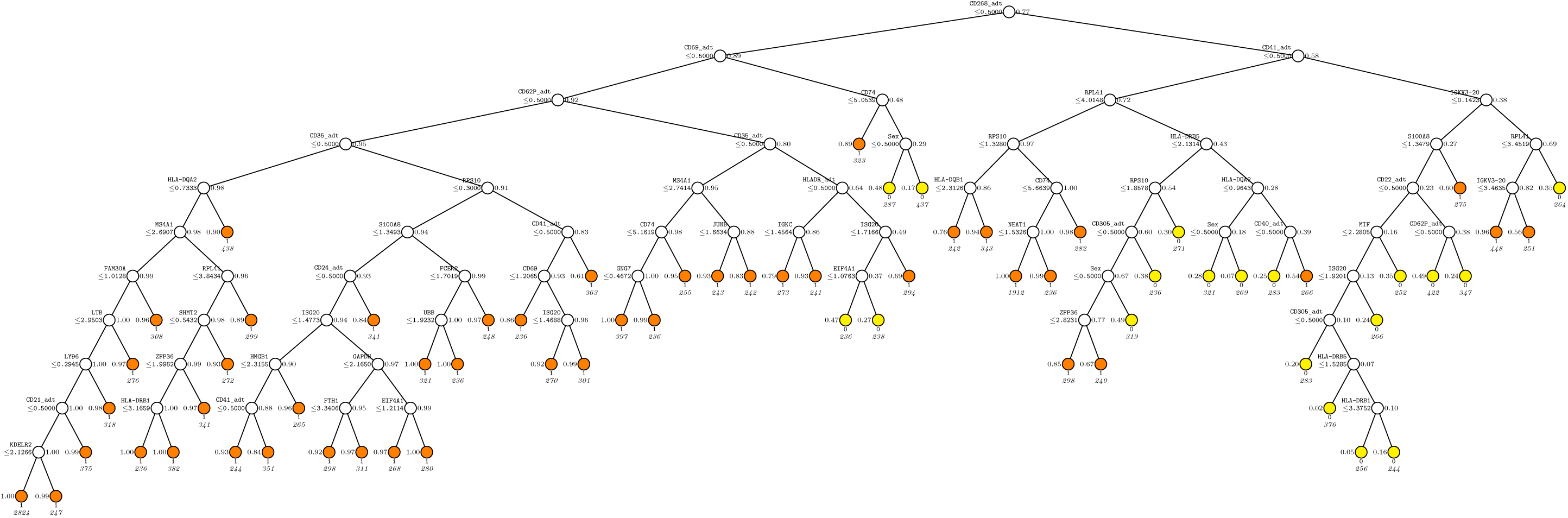
CART tree in B-cell G3. At each split, an observation goes to the left branch if and only if the condition is satisfied. Predicted class and sample size (in italics) printed below each terminal nodes; class sample proportion for Y = 1 beside node. Terminal nodes with proportions of 1s above and below value of 0.5 are painted vermillion and yellow respectively.

**Fig. S18.**
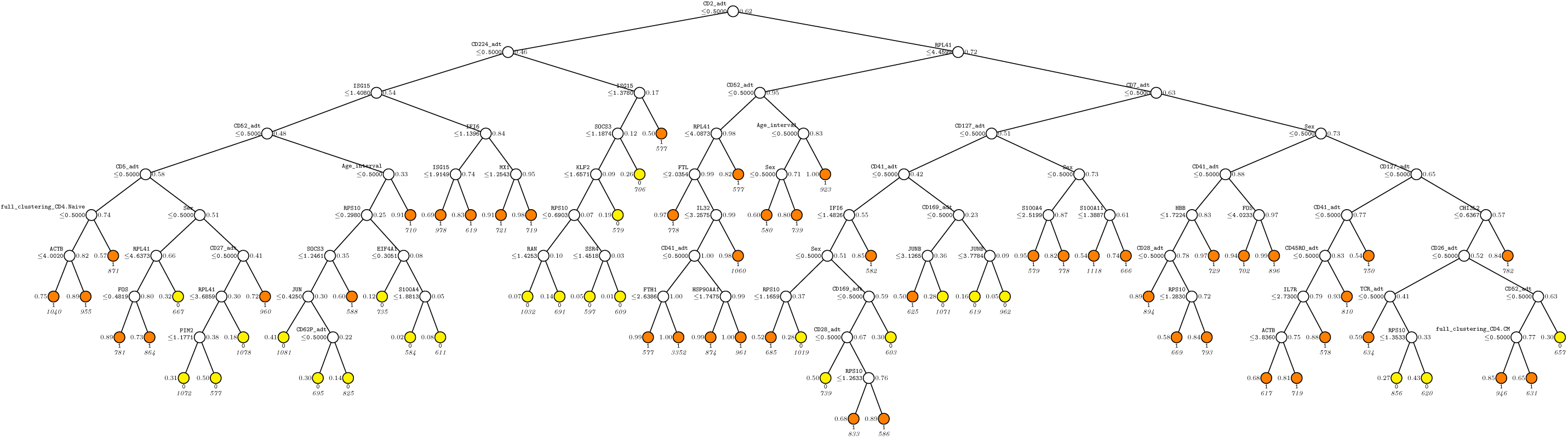
CART tree in CD4T-cell G2. At each split, an observation goes to the left branch if and only if the condition is satisfied. Predicted class and sample size (in italics) printed below each terminal nodes; class sample proportion for Y = 1 beside node. Terminal nodes with proportions of 1s above and below value of 0.5 are painted vermillion and yellow respectively.

**Fig. S19.**
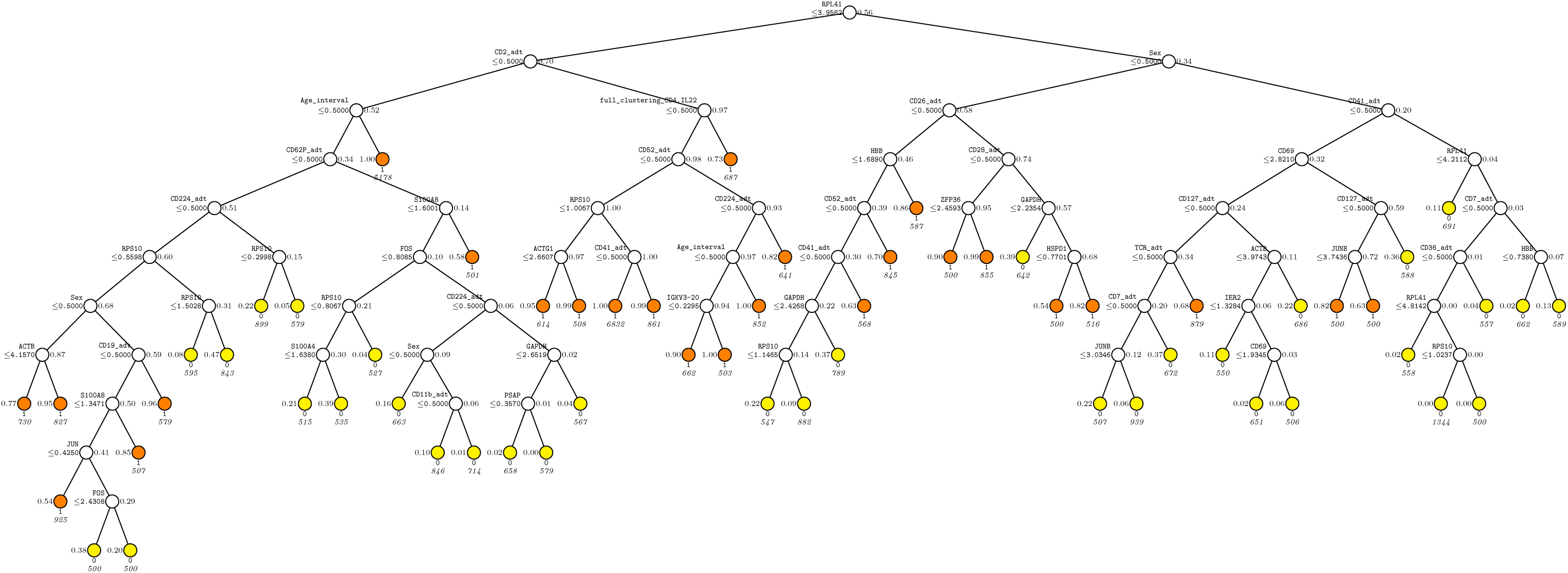
CART tree in CD4T-cell G3. At each split, an observation goes to the left branch if and only if the condition is satisfied. Predicted class and sample size (in italics) printed below each terminal nodes; class sample proportion for Y = 1 beside node. Terminal nodes with proportions of 1s above and below value of 0.5 are painted vermillion and yellow respectively.

**Fig. S20.**
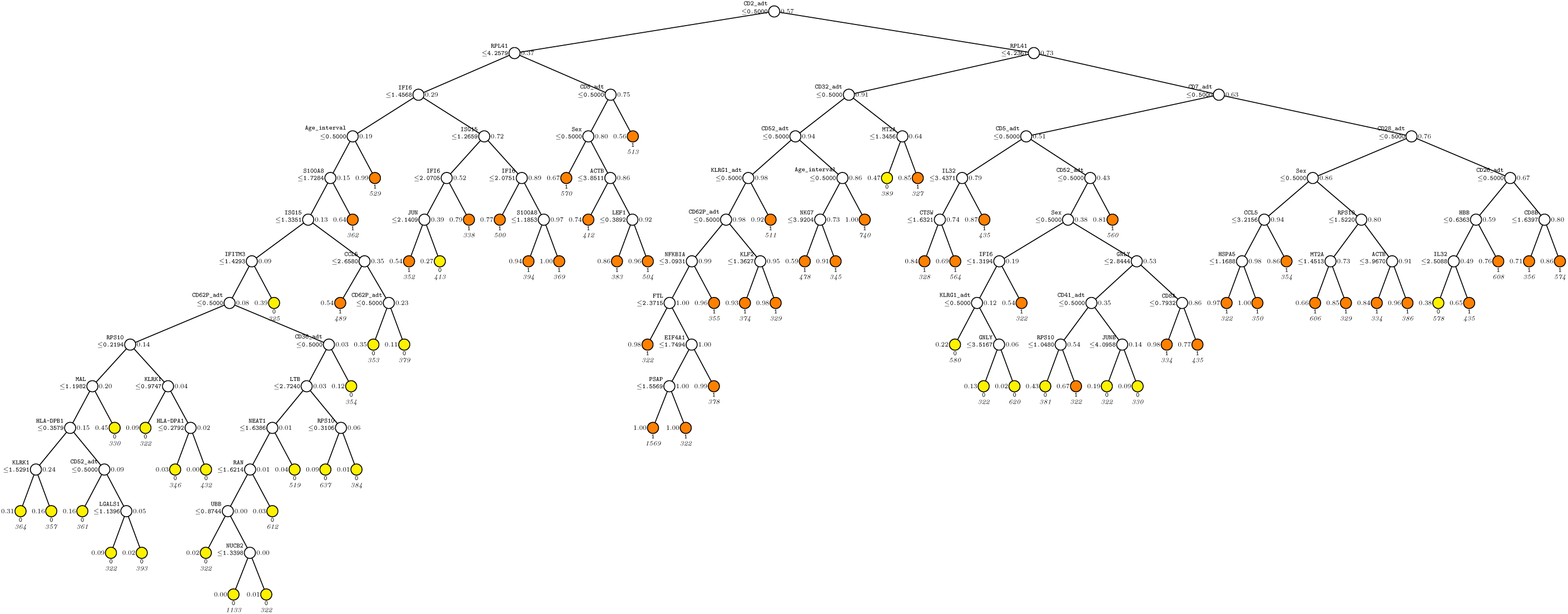
CART tree in CD8T-cell G2. At each split, an observation goes to the left branch if and only if the condition is satisfied. Predicted class and sample size (in italics) printed below each terminal nodes; class sample proportion for Y = 1 beside node. Terminal nodes with proportions of 1s above and below value of 0.5 are painted vermillion and yellow respectively.

**Fig. S21.**
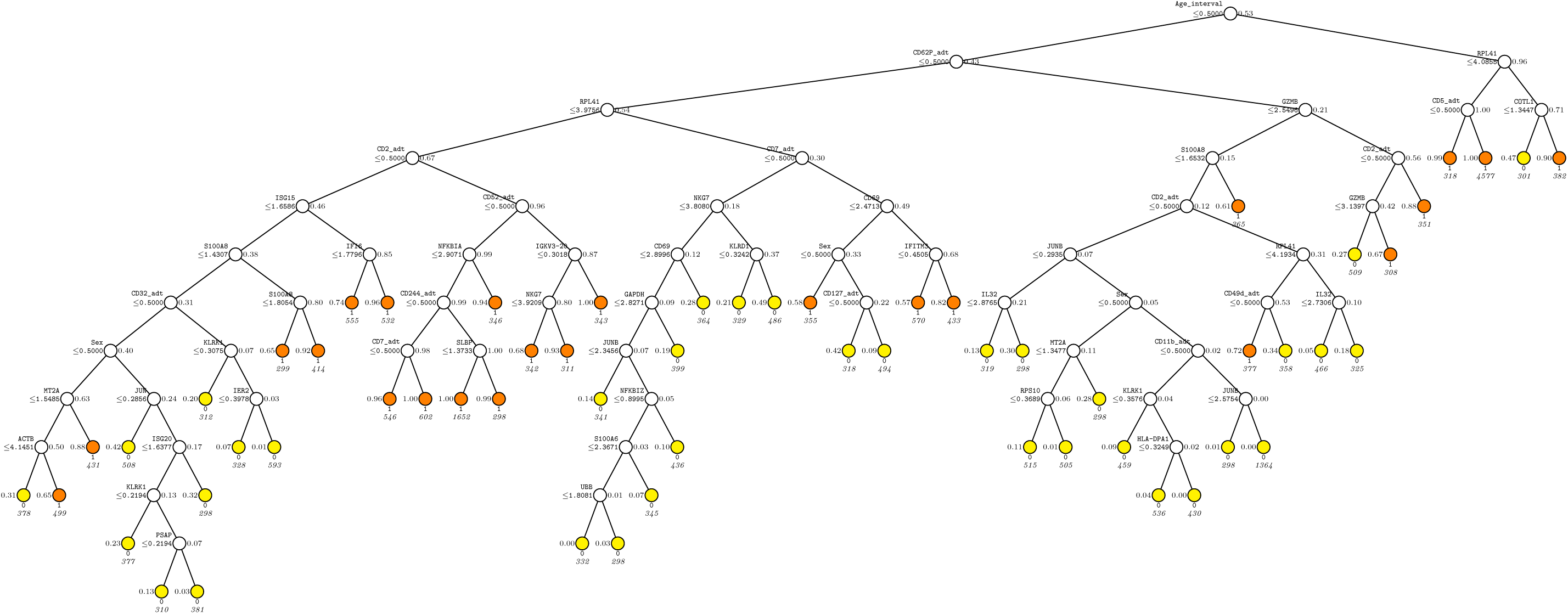
CART tree in CD8T-cell G2. At each split, an observation goes to the left branch if and only if the condition is satisfied. Predicted class and sample size (in italics) printed below each terminal nodes; class sample proportion for Y = 1 beside node. Terminal nodes with proportions of 1s above and below value of 0.5 are painted vermillion and yellow respectively.

**Fig. S22.**
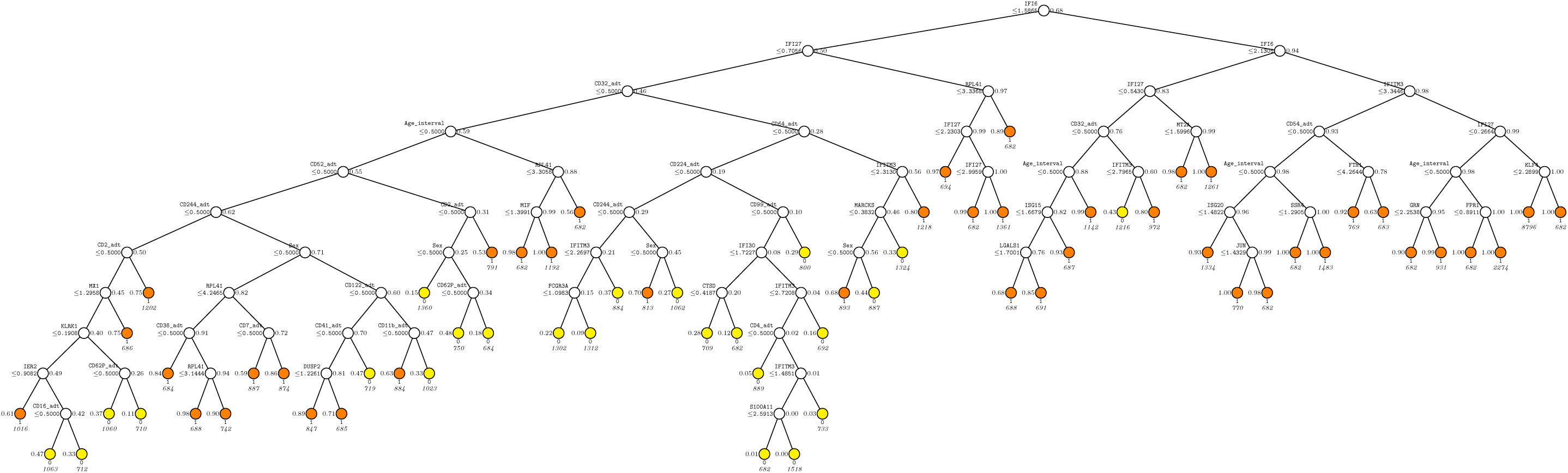
CART tree in Mononuclear-cell G2. At each split, an observation goes to the left branch if and only if the condition is satisfied. Predicted class and sample size (in italics) printed below each terminal nodes; class sample proportion for Y = 1 beside node. Terminal nodes with proportions of 1s above and below value of 0.5 are painted vermillion and yellow respectively.

**Fig. S23.**
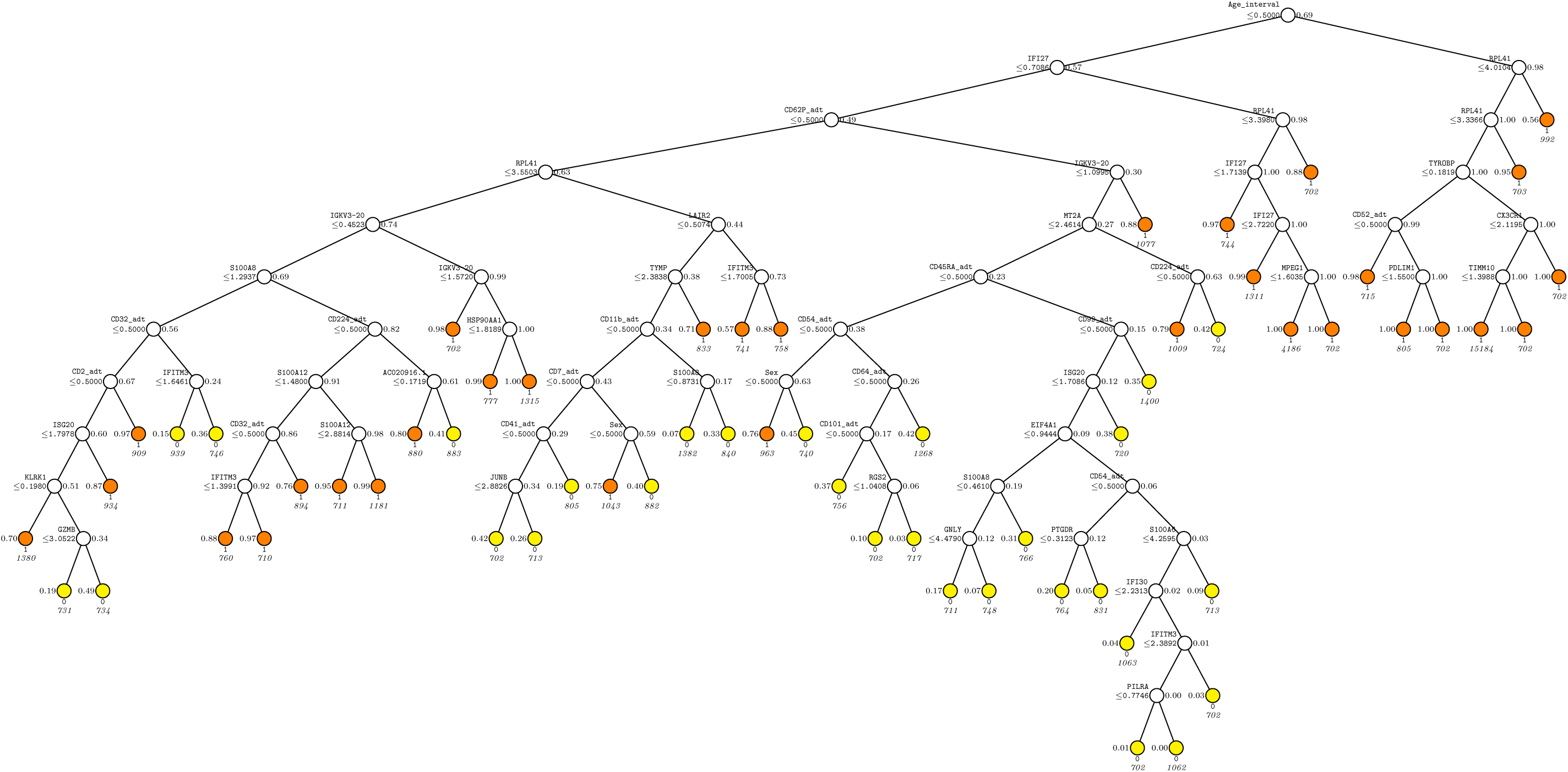
CART tree in Mononuclear-cell G3. At each split, an observation goes to the left branch if and only if the condition is satisfied. Predicted class and sample size (in italics) printed below each terminal nodes; class sample proportion for Y = 1 beside node. Terminal nodes with proportions of 1s above and below value of 0.5 are painted vermillion and yellow respectively.

**Fig. S24.**
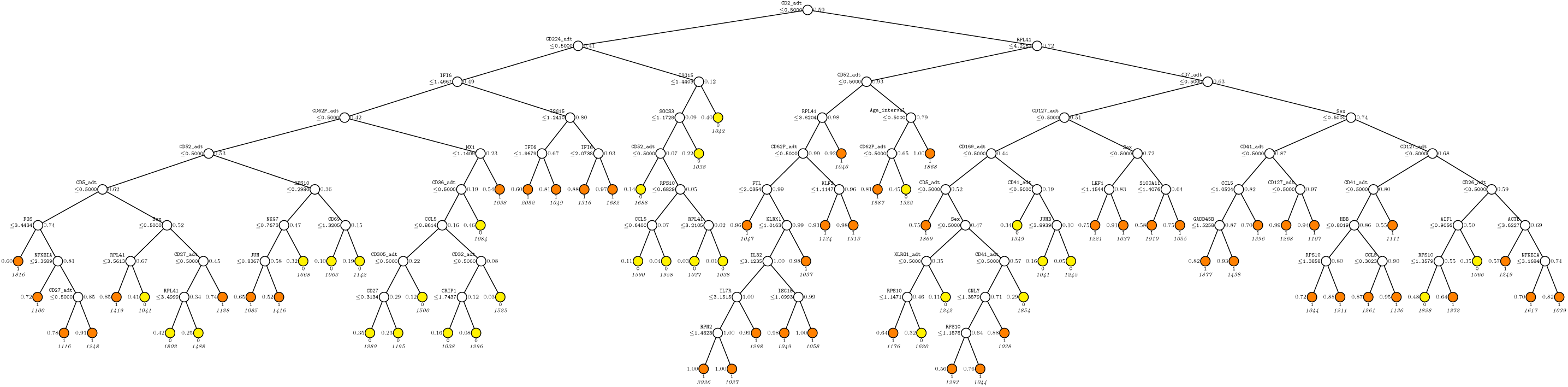
CART tree in T-cell G2. At each split, an observation goes to the left branch if and only if the condition is satisfied. Predicted class and sample size (in italics) printed below each terminal nodes; class sample proportion for Y = 1 beside node. Terminal nodes with proportions of 1s above and below value of 0.5 are painted vermillion and yellow respectively.

**Fig. S25.**
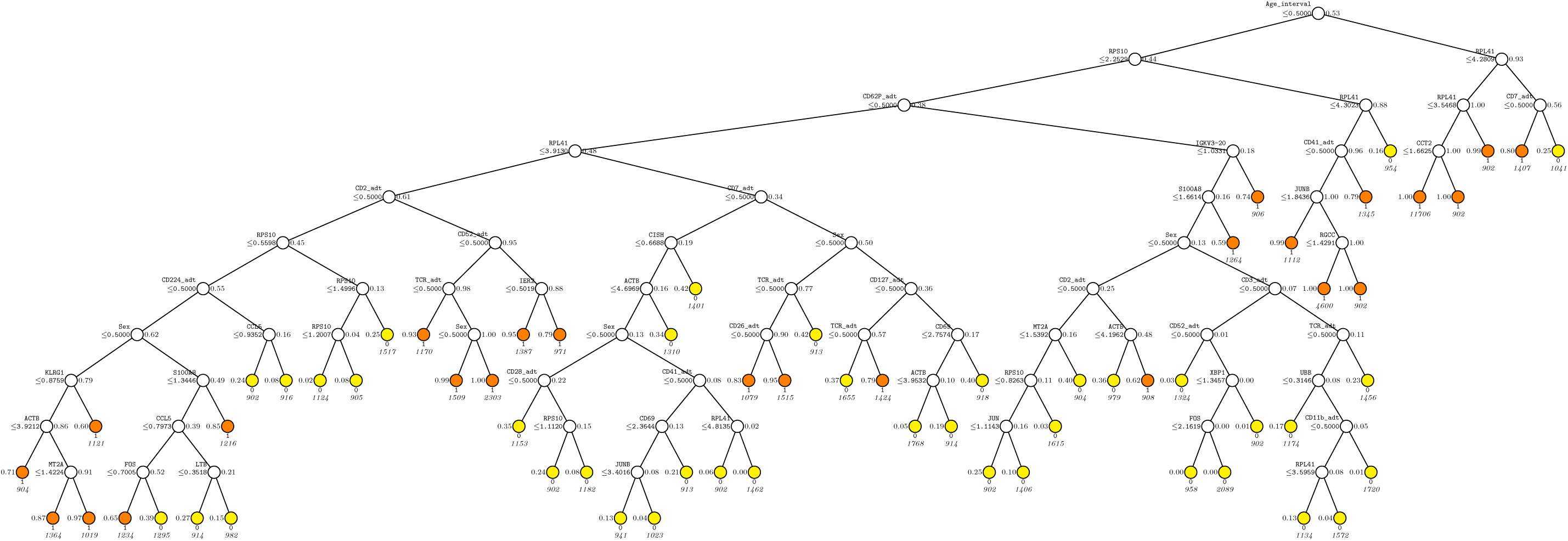
CART tree in T-cell G3. At each split, an observation goes to the left branch if and only if the condition is satisfied. Predicted class and sample size (in italics) printed below each terminal nodes; class sample proportion for Y = 1 beside node. Terminal nodes with proportions of 1s above and below value of 0.5 are painted vermillion and yellow respectively.

